# STX1A localizes to the lysosome and controls its exocytosis

**DOI:** 10.1101/2025.03.29.646068

**Authors:** Anshul Milap Bhatt, Subba Rao Gangi Setty

**Affiliations:** Department of Microbiology and Cell Biology, Indian Institute of Science, Bangalore 560012, India

**Keywords:** STX1A, SNAP23, SNAP25, VAMP2, lysosome exocytosis and SNARE

## Abstract

Lysosome exocytosis is one of the critical functions of lysosomes in maintaining cellular homeostasis and plasma membrane repair. The SNAREs regulating the lysosome fusion with the cell surface at the basal level have been poorly defined. Here, we identified STX1A, a Qa-SNARE localized majorly to lysosomes and a cohort to the plasma membrane in HeLa cells. Overexpression of GFP-STX1A in HeLa cells causes decreased lysosome number and their peripheral dispersion. In line, STX1A knockdown resulted in the accumulation of lysosomes beneath the cell surface and showed reduced lysosome exocytosis in HeLa cells. TIRF imaging microscopy demonstrated an enhanced enrichment of LAMP1-positive vesicles at the cell surface in STX1A depleted compared to control cells. Moreover, STX1A depletion reduces the proteolytic activity without affecting the lysosome content or acidity and increases peripheral lysosome dispersion compared to control cells. Consistently, these cells display an enhanced number of autophagosomes and accumulate autolysosomes. Functionally, GFP-STX1A also localizes to LLOMe-induced GAL3-positive damaged lysosomes and reduces their number by enhancing exocytosis. Biochemically, STX1A forms a SNARE complex with SNAP23 or SNAP25 (Qbc) and VAMP2 (R), and their knockdown in HeLa cells mimics the STX1A- depletion phenotypes. Overall, these studies demonstrate a unique function of STX1A in regulating lysosomal exocytosis by localizing to these degradative organelles.

**Key points:**

- STX1A localizes to lysosomes and plasma membrane in HeLa cells
- STX1A overexpression decreases the lysosome number and their dispersion and facilitates the exocytosis of LLOMe-damaged lysosomes
- STX1A knockdown enhances the lysosome dispersion, accumulates beneath the cell surface in HeLa cells, and displays decreased lysosome exocytosis
- STX1A biochemically interacts with SNAP23/SNAP25, VAMP2 and their individual knockdown phenocopy their phenotypes each other

## Introduction

The essential function of soluble-N-ethylmaleimide-sensitive-factor accessory-protein receptors (SNAREs) in conducting membrane fusion events during intracellular transport has been very well established (Chen and Scheller, 2001; Hong, 2005; Jahn and Scheller, 2006; Sudhof and Rothman, 2009). SNARE family consists of 38 members, and they selectively localize to the specific subcellular membranes and participate in membrane fusion in all cell types (Chen and Scheller, 2001; Hong, 2005). Additionally, SNARE expression and their subcellular localization in different cell types have significantly contributed to the identification and mapping of several intracellular pathways (Chen and Scheller, 2001; Jahn and Scheller, 2006; Wickner and Schekman, 2008). Structurally, SNARE proteins possess a SNARE domain and a C-terminal transmembrane domain or lipid anchor, which will allow them to associate with the membranes. These proteins have been classified into Qa-, Qb-, Qc-, Qbc- (contains two SNARE motifs) and R-SNAREs (Hong, 2005; Jahn et al., 2003). During membrane fusion, two or three specific Q-SNAREs bind to form a cognate partner with R-SNARE on the opposing membrane through their SNARE motifs. This process forms a *trans*-SNARE complex, which brings the two membranes into proximity and promotes the fusion followed by cargo delivery (Sudhof, 2014). Other regulator proteins, such as Sec1/Munc18 (SM) family members, adaptors, and Rabs, modulate the SNARE fusion by controlling their availability and the assembly/disassembly of SNARE complexes (Miller et al., 2007; Stenmark, 2009; Sudhof, 2014; Sudhof and Rothman, 2009).

Syntaxin1A (STX1A) in neurons has been shown to localize to the plasma membrane (PM) (Bennett et al., 1993). STX1A associates with Qbc SNARE synaptosomal-associated protein (SNAP)25 on the PM and then pairs with a presynaptic vesicle R-SNARE vesicle-associated membrane protein (VAMP)2 during the release of neurotransmitters as well as secretion of neuropeptides and neurotrophins (Sudhof, 2014). Further, STX1A has been shown to be enriched at the synapse (Bennett et al., 1992) along with SNAP25 (Oyler et al., 1989). However, earlier immunogold electron microscopy (IEM) studies observed STX1 on the synaptic vesicles (SVs), raising the possibility of SNARE recycling at the nerve terminal (Koh et al., 1993). Consistently, a significant pool (3% of total vesicle protein) of STX1-SNAP25 complex was observed on SVs, possibly contributing to the recycling SNARE pool (Walch-Solimena et al., 1995). PI(3,4,5)P_3_ has been shown to facilitate the clustering of STX1A at the neurotransmitter release sites, indicating a possible role for phosphoinositides during the membrane fusion (Khuong et al., 2013). Interestingly, S^14^ (serine 14) residue phosphorylation in STX1 (phospho-syntaxin 1, pSTX1) occurs during brain development and localizes to the different axonal zones, potentially contributing to the exocytosis (Foletti et al., 2000). STX1A possess SUMOylation sites near the C-terminal transmembrane region, and their prevention affects synaptic vesicle endo/exocytosis balance (Craig et al., 2015), demonstrating the versatility of SNARE function through multiple regulatory mechanisms. Deletion of the N-terminal regulatory domain hampers the localization of STX1A to PM in CHO cells but not INS1 cells (Yang et al., 2006). Additionally, STX1A was shown to bind to granuphilin, an effector of Rab27a, during the insulin excretion in pancreatic cells (Torii et al., 2002). STX1A also participates in secretory functions such as releasing glucagon-like peptide 1 (GLP-1) from intestinal enteroendocrine L cells (Wheeler et al., 2017). Studies have shown that the Munc18a-Syn-1A-SNAP25-VAMP2 complex mediates the exocytosis of pre-docked insulin secretory granules, responsible for the first phase of biphasic glucose-stimulated insulin secretion (GSIS)(Oh et al., 2012; Ohara-Imaizumi et al., 2007). These studies majorly illustrate that STX1A will operate its function from the PM. STX1A has also been found at the endosomes (Brandhorst et al., 2006), and a single isoform was detected on the phagosomes isolated from Drosophila by proteomics approach (Stuart et al., 2007). However, the functional role of STX1 at endosomes and/or phagosomes remains unclear (Dingjan et al., 2018). Thus, studying the intracellular localization of STX1A, including its function in non-neuronal cells, is yet to be comprehensively understood.

Lysosomes play an essential role in many cellular processes, including cargo degradation, gene regulation, immunity, PM repair, cell adhesion and migration (Ballabio and Bonifacino, 2020). Moreover, lysosomes exhibit changes in their dynamics by altering number, distribution, size and fusion/contact with other organelles (de Araujo et al., 2020; Luzio et al., 2007; Yang and Wang, 2021). Lysosomes localize majorly at the perinuclear area and a cohort at the periphery. Studies have shown that various factors regulate lysosome positioning, including cytoskeleton, motor proteins and their contacts with other organelles such as ER, Golgi and mitochondria (Bonifacino and Neefjes, 2017; Cabukusta and Neefjes, 2018; Johnson et al., 2016; Pu et al., 2016). Lysosomes undergo fusion with late endosomes (LE), endosomes and autophagosomes either for cargo delivery or degradation (Ballabio and Bonifacino, 2020). These processes have been shown to be controlled by several SNAREs, including STX7 (Qa), Vti1B (Qb), STX8 (Qc) and VAMP7 (R) mediate the LE-lysosome fusion (Pryor et al., 2004); STX4 (Qa), SNAP23 (Qbc) and VAMP7 (R) regulate the fusion of lysosome with PM (Jaiswal et al., 2002; Rao et al., 2004); and STX17 (Qa), SNAP29 (Qbc) and VAMP7 (Drosophila)/VAMP8 (mammalian) (R) facilitate the autophagosome fusion with lysosomes (Itakura et al., 2012; Takats et al., 2018). Several studies have shown that lysosomes release their content through PM through a process called lysosome exocytosis, which maintains several intra- and extra- cellular processes (Buratta et al., 2020; Samie and Xu, 2014; Tancini et al., 2020; Trojani et al., 2024). During this process, the lysosomes move towards the cell surface and then fuse in a TRPML1-mediated calcium-release manner (Tancini et al., 2020), allowing lysosomal enzymes to be released outside. Lysosome exocytosis has been shown to play a primary role in membrane remodeling and secretion, including PM repair, bone resorption and antigen presentation (Buratta et al., 2020; Samie and Xu, 2014). The role of lysosomal localized VAMP7 (R) and PM localized STX4 (Qa) has been implicated during the fusion of lysosomes with PM (Rao et al., 2004). The role of other SNAREs regulating this process, in addition to its cell type specificity other than the secretory cells, has not been fully understood.

In this study, we attempted to study the intracellular localization and function of STX1A in non-neuronal cell type HeLa and its role in mediating lysosome exocytosis. A variety of biochemical and cell biological approaches demonstrate that GFP-STX1A localizes primarily to lysosomes, and its overexpression in HeLa cells reduces the lysosome number. Depletion of STX1A in HeLa cells enhances the peripheral distribution of lysosomes with a defective PM fusion and reduced enzymatic activity. STX1A localizes to damaged lysosomes induced by LLOMe and facilitates their exocytosis. Mechanically, STX1A uniquely pairs with SNAP23 or SNAP25 on the lysosomes and interacts with VAMP2 on the PM during their exocytosis.

## Results

### STX1A localizes majorly to lysosomes and a cohort to the plasma membrane

STX1A has been reported to localize to the PM in different non-neuronal cell types (Fan et al., 2007; Yang et al., 2006). However, the intracellular localization and function of STX1A in other cell types are poorly defined. Here, we first verified the endogenous expression of STX1A in HeLa cells by immunoblotting (**Fig. S1A**). Due to a lack of commercial antibodies suitable for immunofluorescence microscopy (IFM), STX1A was tagged with GFP at the N-terminus and expressed in the HeLa cells (**Fig. 1A**). The expression of GFP-STX1A was verified using immunoblotting (**Fig. S1B**) and the cells were immunostained for various organelle specific markers to study its intracellular localization. IFM analysis revealed that GFP-STX1A localized as intracellular puncta (**Fig. S1C**) and showed major colocalization with lysosomal proteins LAMP1 (Pearson’s coefficient, *r*=0.64±0.03) and LAMP2 (*r*=0.65±0.02) (**Fig. 1A** and **1B**). In line, GFP-STX1A was also showed higher colocalization with the lysosome peripheral membrane protein Arl8b-tomato (*r*=0.72±0.03), internalized dextran-Alexa fluor (AF) 594 (*r*=0.55±0.02), acidic organelle marker lysoTracker-Red (*r*=0.52±0.02) and DQ-Red-BSA positive active lysosomes (*r*=0.49±0.02) (**Fig. 1A** and **1B**). Interestingly, a cohort of GFP- STX1A was observed in late endosomes marked by LBPA (*r*=0.51±0.02) (**Fig. 1A** and **1B**). However, GFP-STX1A did not exhibit its colocalization to early endosomes (EEA1) (*r*=0.18±0.02), *trans*-Golgi (p230) (*r*=0.14±0.01) or *cis*-Golgi (GM130) (*r*=0.16±0.02) (**Fig. 1A** and **1B**). Live-cell imaging of HeLa cells showed that GFP-STX1A colocalizes completely with LAMP1-RFP positive lysosomes and moves together throughout the time frame (**Fig. S1D**). Further, mCherry-STX1A made contact with GFP-CD63 marked compartments, similar to LAMP1-RFP with GFP-CD63 (**Fig. S1E** and **S1F**). These studies demonstrate the intracellular localization of STX1A to lysosomes in HeLa cells. As expected, a cohort of GFP-STX1A was also observed on the cell surface and showed colocalization with the PM, marked by wheat germ agglutinin (WGA)-AF 594 (**Fig. 1C**). Furthermore, GFP-STX1A was observed in the vicinity of the cortical actin marked by phalloidin AF 594 or mCherry-UtrCH to indicates its localization to the PM (**Fig. 1C**). However, sucrose gradient membrane fractionation studies showed endogenous STX1A participates more to the LAMP1-positive membrane fractions compared to the cell surface (labelled with Na^+^-K^+^ ATPase) or to the recycling endosomes (labelled with Rab22A) (**Fig. S1G**), indicating a unique localization of STX1A to the lysosomes.

**Figure 1.**
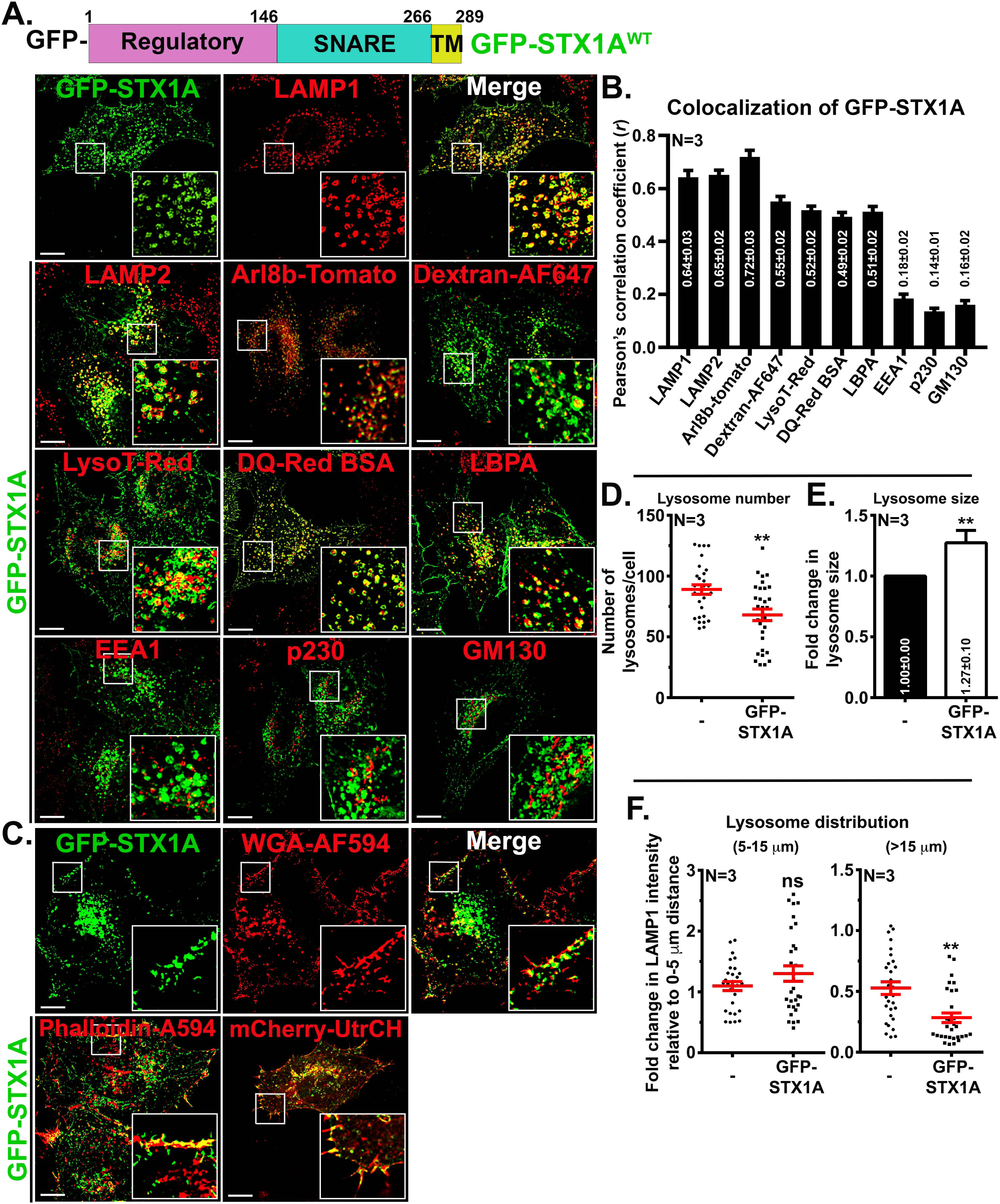
STX1A localizes majorly to lysosomes and plasma membrane. (A) Schematic representation of STX1A. TM, transmembrane domain. Domain lengths are indicated by amino acid numbers. (A, C) IFM analysis of HeLa cells expressing GFP-STX1A. Cells were stained with antibodies (anti-LAMP1, anti-LAMP2, anti-LBPA, anti-EEA1, anti-p230, anti-GM130) or subjected to uptake (Dextran-AF647, LysoTracker-Red or DQ-Red BSA), stained (WGA-AF594 or phalloidin-AF594) or transfected with Arl8b-tomato or mCherry-UtrCH. Insets are magnified views of the white boxed areas. Scale bars, 10 µm. (B) Plot represents the Pearson’s correlation coefficient (*r*) of GFP-STX1A with various organelle markers as listed. (D) Plot represents the number of lysosomes/cell in the non-transfected (-) or GFP-STX1A expressing cells. (E) Plot represents the fold change in lysosome size in the non-transfected (-) or GFP-STX1A expressing cells. (F) Plots represent the ratio of change in radial profile intensity (represents lysosome dispersion) of lysosomes between 5-15 µm or more than 15 µm with respect to 0-5 µm, respectively. The average values in mean±s.e.m. are indicated on the graphs. N=3. ns, non-significant and **p≤0.01.

To study the effect of GFP-STX1A overexpression on lysosomes, we compared non-transfected with the transfected HeLa cells and measured the lysosome number, activity, acidity, size and distribution in the cells. Interestingly, the average lysosome number was significantly reduced in cells overexpressing GFP-STX1A compared to non-transfected cells (non-transfected cells: 89±4 and GFP-STX1A cells: 68±5) (**Fig. 1D**). However, the number of lysoTracker-Red positive compartments (non-transfected cells: 241±10 and GFP-STX1A cells: 223±17) and active lysosomes positive for DQ-Red-BSA (measured as corrected total cell fluorescence intensity, CTCF for non-transfected cells: 2.47±0.22x10^6^ A.U. and GFP-STX1A cells: 1.91±0.31x10^6^ A.U.) were not altered upon transfection of cells with GFP-STX1A (**Fig. S1H** and **S1I**). Unexpectedly, the size of lysosomes was significantly enhanced upon overexpression of GFP- STX1A compared to non-transfected cells (non-transfected cells: 0.35±0.02 μm^2^ and GFP- STX1A cells: 0.42±0.02 μm^2^) (**Fig. 1E** and **S1J**). Furthermore, the fold change in the distribution of lysosomes (measured by radial profile intensity measurement) between 5-15 μm distance was unaffected with respect to 0-5 μm distance in the cells (non-transfected cells: 1.10±0.07 and GFP-STX1A cells: 1.30±0.13); however, it was significantly reduced beyond >15 μm distance (non-transfected cells: 0.53±0.05 and GFP-STX1A cells: 0.28±0.04) (**Fig. 1F**). Overall, these data demonstrate that GFP-STX1A majorly localizes to the lysosomes and a cohort to the PM. STX1A overexpression increases lysosome size and reduces lysosome number and peripheral dispersion.

### Expression of regulatory domain deletion mutant, Rab7 depletion or lysosomal damage in HeLa cells affects the localization of STX1A to lysosomes

The N-terminus of the syntaxin family SNAREs serves as a regulatory domain in determining the specificity of fusion and localization (Shen et al., 2010). To investigate the role of regulatory domain in STX1A, we deleted the amino acids from 25 to 146 in GFP-STX1A using site-directed mutagenesis (GFP-STX1A^Δ25-146^) (**Fig. 2A**). IFM studies showed that GFP-STX1A^Δ25-146^ mutant colocalized marginally with LAMP1-positive lysosomes (*r*=0.20±0.03) compared to the wildtype GFP-STX1A (*r*=0.56±0.02) (**Fig. 2B** and **2C**). A cohort of the GFP-STX1A^Δ25-146^ mutant was observed in the Golgi, as indicated by its colocalization with p230 (*r*=0.24±0.04) compared to GFP-STX1A (*r*=0.09±0.02) (**Fig. 2B** and **2C**) in addition to the cell surface (refer **Fig. S5B**). Next, we treated the cells with nocodazole to disperse the lysosomes (LAMP1), which showed no major colocalization with GFP-STX1A^Δ25-146^, whereas GFP-STX1A showed an overlap with LAMP1 positive lysosomes (**Fig. S2A**). Overexpression of GFP-STX1A^Δ25-146^ mutant in HeLa cells showed decreased DQ-Red BSA activity compared to GFP-STX1A^WT^ cells (CTCF for non-transfected cells: 2.47±0.22x10^6^ A.U. and GFP-STX1A^Δ25-146^ cells: 1.15±0.18x10^6^ A.U.) (**Fig. S2B** and **S2C**). Consistently, the number of DQ-Red puncta was significantly reduced in cells expressing GFP-STX1A^Δ25-146^ compared to non-expressing cells (**Fig. S2B** and **S2C**). However, the number of lysotracker puncta did not significantly alter upon expression of GFP-STX1A^Δ25-146^ mutant in HeLa cells (non-transfected cells: 241±11 and GFP- STX1A^Δ25-146^ cells: 279±30) (**Fig. S2D** and **S2E**). Interestingly, the total number of lysosomes was dramatically decreased upon expression of GFP-STX1A^Δ25-146^ mutant (**Fig. 2D**) (non-transfected cells: 89±4 and GFP-STX1A^Δ25-146^ cells: 63±6). In contrast, the size of LAMP1 positive lysosomes was enhanced with the expression of GFP-STX1A^Δ25-146^ mutant compared to non-expressing cells (**Fig. 2E**) (non-transfected cells: 0.35±0.03 μm^2^ and GFP-STX1A^Δ25-146^: 0.70±0.07 μm^2^). Further, the fold change in the lysosome size upon GFP-STX1A^Δ25-146^ (2.37±0.29 folds) overexpression is significantly higher in comparison to non-expressing cells (**Fig. 2E**) or GFP-STX1A expressing cells (1.27±0.10 folds of non-expressing cells) (**Fig. 1E**). In line to the lysosome number, the distribution of LAMP1-positive compartments was significantly reduced at periphery (above 15 µm distance) with the expression of GFP-STX1A^Δ25-146^ mutant compared to non-expressing cells (**Fig. 2F**) (non-transfected cells: 1.11±0.10 for 5-15 µm, 0.57±0.07 for above 15 µm and GFP-STX1A^Δ25-146^: 1.07±0.13 for 5-15 µm, 0.24±0.04 for above 15 µm). These studies illustrate the role of a regulatory domain in the trafficking of STX1A to lysosomes.

**Figure 2.**
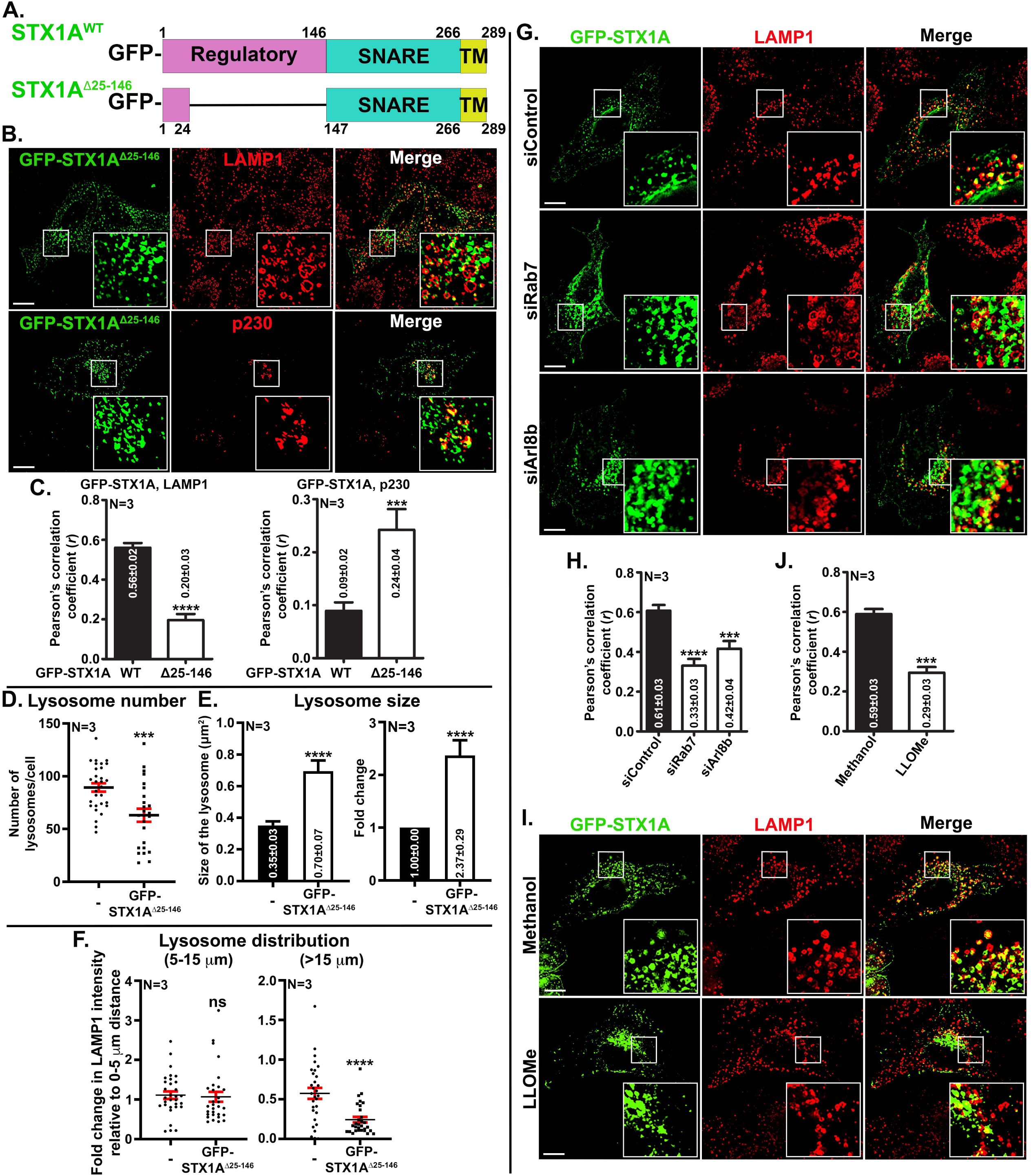
N-terminal regulatory domain of STX1A, Rab7 and lysosomal damage affect the localization of STX1A to lysosomes. (A) Schematic representation of GFP-STX1A and GFP-STX1A^Δ25-146^. TM, transmembrane domain. Domain lengths are indicated by amino acid numbers. (B) IFM analysis of HeLa cells expressing GFP-STX1A^Δ25-146^ and stained for LAMP1 (lysosomes) or p230 (Golgi). (C) Plots represent the Pearson’s correlation coefficient (*r*) of GFP-STX1A and GFP-STX1A^Δ25-146^ with LAMP1 or p230 separately. (D) Plot represents the number of LAMP1 puncta/cell in the non-transfected (-) or GFP-STX1A^Δ25-146^ expressing cells. (E) Plots represent the size of LAMP1 puncta (μm^2^) and their fold difference in the non-transfected (-) or GFP-STX1A^Δ25-146^ expressing cells. (F) Plots represent the ratio of change in radial profile intensity of lysosomes between 5-15 µm distance or more than 15 µm distance with respect to 0-5 µm distance, respectively. (G) IFM analysis of HeLa cells transfected with control siRNA, Rab7 siRNA and Arl8b siRNA. Post 24 h, the cells were again transfected with GFP-STX1A, fixed after 24 h and stained with an anti-LAMP1 antibody. (H) Plot represents the Pearson’s correlation coefficient (*r*) of GFP-STX1A with LAMP1 in siControl, siRab7 and siArl8b cells. (I) HeLa cells were transfected with GFP-STX1A. After 24 h of transfection, cells were treated with either methanol or 3 mM LLOMe for 4 h followed by fixation. The cells were stained for LAMP1 and then analyzed by IFM. Insets in IFM images are magnified views of the white boxed areas. Scale bars, 10 µm. (J) Plot represents the Pearson’s correlation coefficient (*r*) of GFP-STX1A with LAMP1 in methanol or LLOMe treated cells. The average values in mean±s.e.m. are indicated on the graphs. N=3. ns, non-significant, ***p≤0.001 and ****p≤0.0001.

Next, we tested the role of Rab7 and Arl8b in the trafficking of STX1A to lysosomes. Co-expression of mCherry-STX1A with GFP-Rab7^WT^ (*r*=0.58±0.10), GFP-Rab7^Q67L^ (constitutive active form, *r*=0.49±0.07), or GFP-Rab7^T22N^ revealed that the dominant negative mutant expression resulted in diffused distribution of STX1A in the cytosol and was not targeted to LAMP1 compartments (**Fig. S2F**). To further validate these results, we tested the localization of STX1A in Rab7 or Arl8b knockdown cells. The knockdown of Rab7 in HeLa cells led to a decreased colocalization of GFP-STX1A to the lysosomes (siControl: *r*=0.61±0.03, siRab7: *r*=0.33±0.03 and siArl8b: *r*=0.42±0.04) (**Fig. 2G** and **2H**). However, the STX1A protein levels were unaffected in Rab7 knockdown cells compared to control (**Fig. S2G**). The knockdown of Arl8b in HeLa cells also led to a slight decrease in targeting STX1A to lysosomes (**Fig. 2G** and **2H**). This may be due to reduced levels of Rab7 in the Arl8b knockdown cells (**Fig. S2G**). These studies illustrate that Rab7 and/or Arl8 regulate STX1A trafficking to the lysosomes.

LLOMe has been shown to cause damage to lysosomes and other endocytic organelles (Eriksson et al., 2020). Here, we tested whether the damage of lysosomes perturb the trafficking of STX1A to lysosomes. Cells expressing GFP-STX1A were treated with methanol or LLOMe and studied its localization to lysosomes (**Fig. 2I**). Interestingly, GFP-STX1A showed a decreased localization to the LAMP1-positive compartments (Methanol: *r*=0.59±0.03, LLOMe: *r*=0.29±0.03) (**Fig. 2I** and **2J**). In contrast, mCherry-STX1A localizes to the damaged lysosomes marked by YFP-GAL3, unlike LAMP1-positive compartments (**Fig. S2H**). Additionally, we noted that the number of damaged lysosomes marked by YFP-GAL3 also significantly reduced in the mCherry-STX1A expressing cells as compared to non-expressing cells (0.64±0.08 folds) (**Fig. S2H** and **S2I**). In line, the YFP-GAL3-positive structures were moderately positive for LAMP1 in LLOMe treated cells (**Fig. S2H**) compared to methanol treated cells. In conclusion, these results indicate that the regulatory domain of STX1A plays a role in its localization to the lysosomes. The trafficking of STX1A appeared to be dependent on the GTPase activity of Rab7. Lastly, the lysosome damage by LLOMe affects the trafficking of STX1A to LAMP1-positive lysosomes, and the overexpression of mCherry-STX1A in HeLa cells reduces the number of damaged lysosomes. Overall, these studies demonstrate a role for STX1A in regulating lysosome function.

### Knockdown of STX1A in HeLa cells leads to an increase in lysosome dispersion and a decrease in lysosome exocytosis

To examine the function of lysosomal localized STX1A in HeLa cells, we depleted the SNARE using two different shRNAs or complementary siRNAs (referred to here as sh/siSTX1A-1/-2 and sh/siControl) and confirmed the knockdown by immunoblotting. HeLa cells showed more than 50% loss of STX1A protein with two different target sequences compared to control sh/siRNA (**Figs. S3A** and **S3B**). The STX1A knockdown cells were further immunostained for LAMP1 to mark the lysosomes and with phalloidin AF 488, which marks the cell boundary. (**Fig. 3A**, **Figs. S3A** and **S3B**). Interestingly, the lysosomes displayed enhanced distribution in STX1A knockdown compared to control cells (**Fig. 3A**), and this phenotype is consistent with shRNA (sh) or siRNA (si) mediated knockdown of STX1A in HeLa cells (**Fig. S3A** and **S3B**). In line, the measurement of lysosome distribution in the STX1A knockdown cells showed a significant increase in peripheral distribution beyond 15 μm distance (shControl: 0.22±0.02, shSTX1A-1: 0.43±0.03, and shSTX1A-2: 0.40±0.03) compared to 5-15 μm distance from the nucleus (shControl: 1.00±0.07, shSTX1A-1: 1.25±0.08, and shSTX1A-2: 1.06±0.05) (**Fig. 3B**). To confirm the specificity of this phenotype, we used 3’-UTR specific siRNA to knockdown STX1A in HeLa cells and then rescued with GFP-STX1A overexpression (**Fig. S3C** and **S3D**). IFM studies of rescued STX1A knockdown cells restored the lysosomal distribution and appeared similar to siControl cells (**Fig. S3D**). In line, quantification of lysosomal distribution in the rescued cells restored to the siControl cells (at >15 μm distance) (**Fig. S3E**) (non-transfected cells: shControl - 0.26±0.02, shSTX1A-3’UTR - 0.49±0.05, and GFP-STX1A cells: shControl - 0.20±0.02, shSTX1A-3’UTR - 0.27±0.03). These experiments confirm the specificity of si/shRNA used for the depletion of STX1A in the HeLa cells. Further, these studies illustrate that STX1A depletion enhances the lysosome dispersion in HeLa cells.

**Figure 3.**
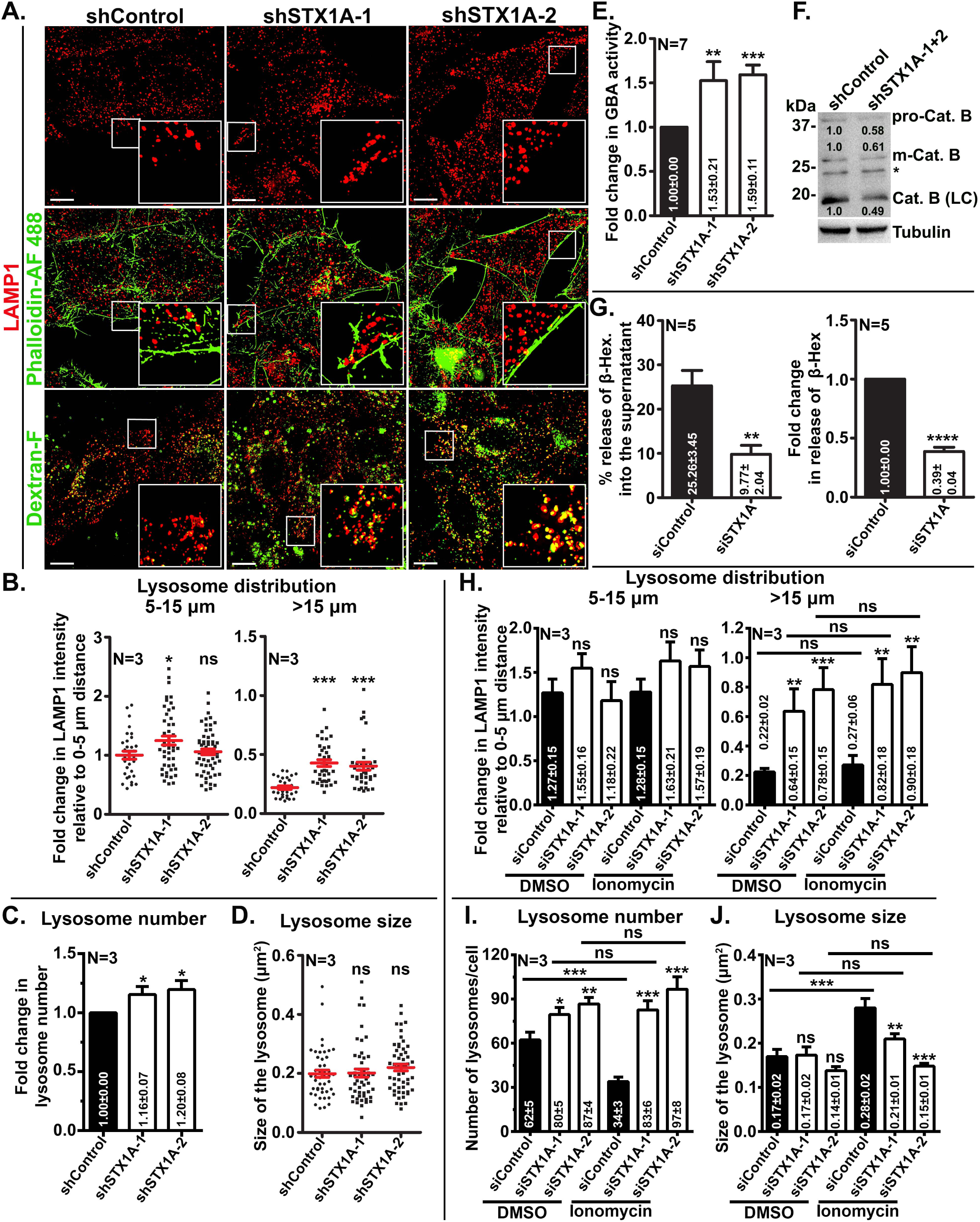
STX1A depletion in HeLa cells leads to an increase in lysosome dispersion and a decrease in lysosome exocytosis. (A) IFM analysis of HeLa cells transduced with lentivirus containing control shRNA (shControl) or two different shRNAs against STX1A (shSTX1A-1 or shSTX1A-2). Cells were stained with anti-LAMP1 antibody and phalloidin-Alexa Fluor 488 or subjected to the uptake of dextran-Fluorescein (Dextran-F). Individual and merged panels are shown separately. Insets are magnified views of the white boxed areas. Scale bars, 10 µm. (B) Plots represent the ratio of change in radial profile intensity of lysosomes between 5-15 µm or more than 15 µm distance with respect to 0-5 µm distance, respectively, as a measure of lysosome dispersion. (C, D, E) Plots represent the fold change in lysosome number (C), size of the lysosomes (μm^2^) (D) and fold change in GBA activity (F) in control and STX1A knockdown cells. (F) Immunoblotting of cell lysates probed for cathepsin B processivity. Pro- immature, m- mature, and LC- light chain. * non-specific band highlighted by anti-cathepsin B antibody. γ- tubulin is used as the loading control. The fold change in band intensities was normalized with internal control and indicated on the blots. (G) Plot represents the percentage change in the release of β-hexosaminidase (β-Hex) into the supernatant as a measure of lysosome exocytosis. Similarly, fold change in the release of β-Hex into the supernatant is shown separately. (H) Plots represent the ratio of change in radial profile intensity of lysosomes between 5-15 µm or more than 15 µm distance with respective to 0-5 µm distance, respectively. Cells were either treated with DMSO or ionomycin. (I, J) Plots represent the number of lysosomes/cell (I) and the size of lysosomes (μm^2^) (J) in the given conditions. The average values in mean±s.e.m. are indicated on the graphs. N=3. ns- nonspecific; *p≤0.1, **p≤0.01; ***p≤0.001 and ****p≤0.0001.

To characterize the functionality of lysosomes, STX1A knockdown cells were internalized with dextran-Fluorescein (dextran-F) to label the terminal lysosomes. IFM studies showed that peripherally distributed lysosomes (LAMP1-postive) in STX1A depleted cells were positive for dextran-F (**Fig. 3A**). Further, quantification of lysosome dynamics in STX1A knockdown cells showed slightly enhanced lysosome number (in fold change - shControl: 1.00±0.00, shSTX1A-1: 1.16±0.07, and shSTX1A-2: 1.20±0.08) with no change in lysosome size compared to control cells (shControl: 0.20±0.01 μm^2^, shSTX1A-1: 0.20±0.01 μm^2^, and shSTX1A-2: 0.22±0.01 μm^2^) (**Figs. 3C** and **3D**). Several studies have shown that peripheral lysosomes possess less enzyme activity due to reduced acidity than the perinuclear localized lysosomes (Korolchuk et al., 2011). Therefore, we tested the lysosomal acidity with LysoTracker Red and proteolytic processivity with the DQ-Red BSA probe in the cells. Interestingly, STX1A depleted cells showed no significant changes in LysoTracker Red staining compared to control cells (shControl: 1.00±0.00, shSTX1A-1: 1.17±0.10, and shSTX1A-2: 0.90±0.07) (**Fig. S4A**). However, STX1A depleted cells displayed a significant decrease in the fluorescence intensity (measured as CTCF) of DQ-Red-positive organelles compared to control cells (shControl: 2.38±0.18x10^6^ A.U., shSTX1A-1: 1.72±0.12x10^6^ A.U., and shSTX1A-2: 1.64±0.13x10^6^ A.U.) (**Fig. S4A**). Next, we tested the lipid hydrolase activity in the lysosomes by measuring β-glucocerebrosidase (β-GC, *GBA1*) activity (Patel et al., 2023b). Surprisingly, STX1A knockdown cells showed significantly enhanced GBA activity (shControl: 1.00±0.00, shSTX1A-1: 1.53±0.21, and shSTX1A-2: 1.59±0.11) (**Fig. 3E**) with a concomitant increase in protein expression (refer **Fig. 4C**) compared to control cells, indicating the enhanced activity is possibly due to enhanced expression. In contrast, the levels of cathepsin B protease are moderately reduced in STX1A depleted compared to control cells (**Fig. 3F**). Overall, these studies suggest that STX1A depletion causes the altered lysosomal number, distribution, and activity without affecting their size and acidity.

**Figure 4.**
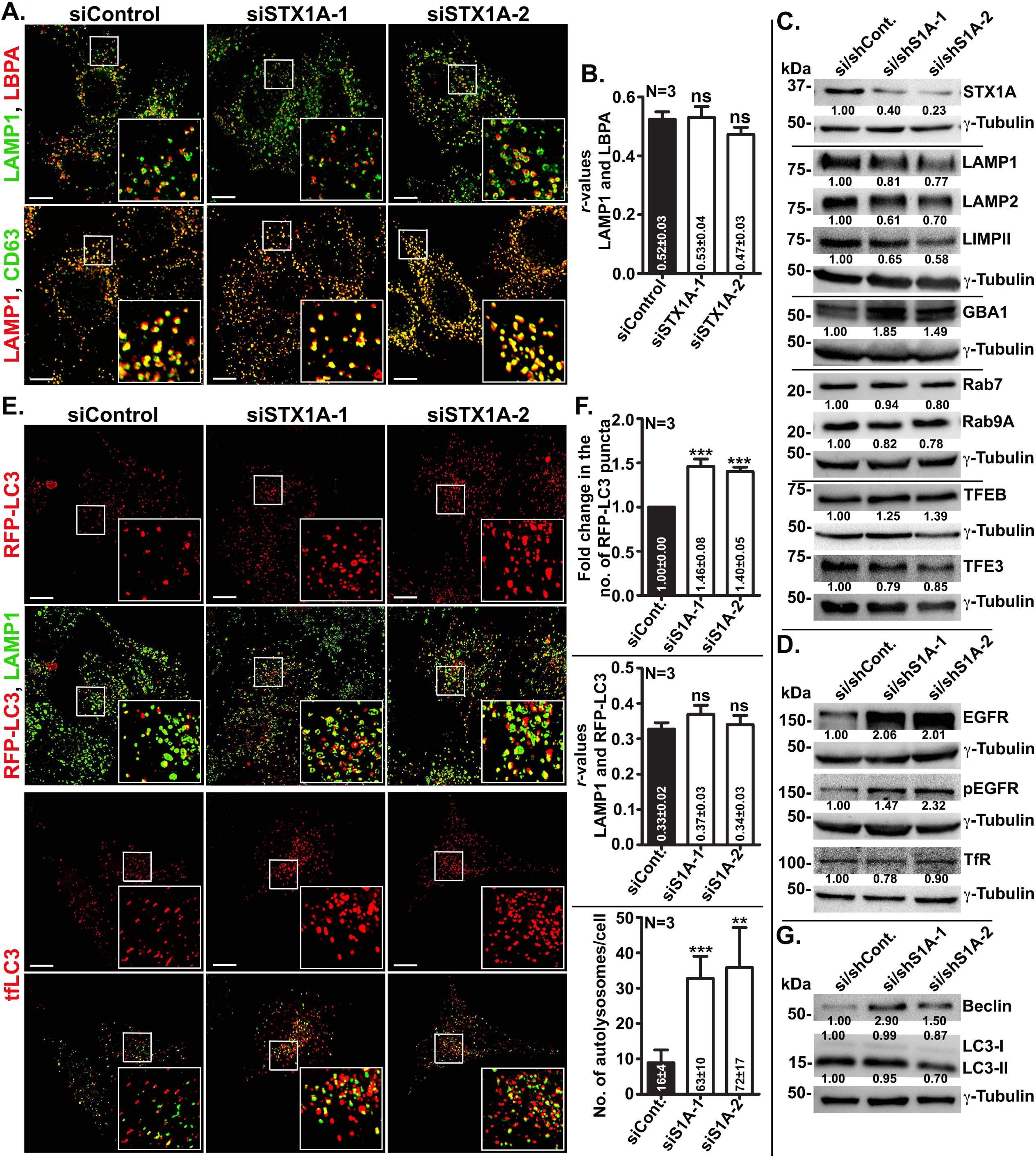
STX1A depletion in HeLa cells displays decreased lysosome degradative capacity and increased autolysosome number. (A, E) IFM analysis of HeLa cells transfected with control or two different STX1A (1 and 2) siRNA sequences. The cells were stained for LAMP1 and LBPA or CD63 (A). Another set of cells was overexpressed with RFP-LC3 or tfLC3 (rat LC3 fused to mRFP and EGFP, E). RFP-LC3 expressing cells were stained for LAMP1. Individual and merged panels are shown separately. Insets are magnified views of the white boxed areas. Scale bars, 10 µm. (B) Plot represents the Pearson’s correlation coefficient (*r*) between LAMP1 and LBPA, shown in A. (C, D, G) Immunoblotting analysis of control and STX1A knockdown cell lysates. The blots were probed for checking the expression of different lysosome associated proteins in C, cargo proteins in D, and autophagy related proteins in G. γ- tubulin is used as the loading control in all immunoblots. The fold change in band intensities was normalized with internal control and indicated on the blots. (F) Plots represent the number of RFP-LC3 puncta, Pearson’s correlation coefficient (*r* value) between LAMP1 and RFP-LC3, and the number of autolysosomes. The average values in mean±s.e.m. are indicated on the graphs. N=3. ns-non-significant, **p≤0.01, and ***p≤0.001.

The depletion of STX1A in HeLa cells led to the observation that the lysosomes were not only dispersed but enhanced their localization beneath the cell surface (**Figs. 3A** and **S2A**, **S2B**). The accumulation of lysosomes nearer to the cell surface in STX1A depleted cells is possibly due to a defect in their exocytosis. To confirm this hypothesis, we performed ionomycin-induced lysosome exocytosis (Rodriguez et al., 1997) in the cells. Ionomycin is known to trigger lysosome exocytosis, which can be evaluated by measuring the percentage of β-hexosaminidase (β-Hex) released into the supernatant (Escrevente et al., 2021). As expected, the percentage of β- Hex released into the supernatant of cells depleted with STX1A showed a significant reduction (0.39±0.04 folds) compared to control cells, indicating a possible role for STX1A in lysosome exocytosis (siControl: 25.26±3.45% and siSTX1A: 9.77±2.04%) (**Fig. 3G**). To further confirm these results, we stained the cells with LAMP1 to label the lysosomes and phalloidin to mark the cell boundary post-ionomycin treatment (**Fig. S4B**) and then measured the lysosome dispersion. In line, the fold distribution of LAMP1-positive lysosomes beyond the 15 µm distance is significantly increased compared to 5-15 µm distance in reference to 0-5 µm distance (**Fig. 3H** and **S4B**) (for 5-15 µm distance: siControl-DMSO: 1.27±0.15, siSTX1A-1-DMSO: 1.55±0.16, siSTX1A-2-DMSO: 1.18±0.22, siControl-ionomycin: 1.28±0.15, siSTX1A-1-ionomycin: 1.63±0.21, siSTX1A-2-ionomycin: 1.57±0.19) (for above 15 µm: siControl-DMSO: 0.22±0.02, siSTX1A-1-DMSO: 0.64±0.15, siSTX1A-2-DMSO: 0.78±0.15, siControl-ionomycin: 0.27±0.06, siSTX1A-1-ionomycin: 0.82±0.18, siSTX1A-2-ionomycin: 0.90±0.18). Additionally, the ionomycin treated control cells showed a significant reduction in the total number of lysosomes as compared to DMSO treated control cells (**Fig. 3I** and **S4B**), proving the evidence that the lysosome exocytosis assay is working. In contrast to control cells, the lysosome number was not altered in STX1A depleted cells treated with ionomycin compared to DMSO treatment (siControl-DMSO: 62±5, siSTX1A-1-DMSO: 80±5, siSTX1A-2-DMSO: 87±4, siControl-ionomycin: 34±3, siSTX1A-1-ionomycin: 83±6, siSTX1A-2-ionomycin: 97±8) (**Fig. 3I**). Moreover, the ionomycin treated STX1A knockdown cells possess significantly enhanced number of lysosomes compared to ionomycin treated control cells (siSTX1A-1: 2.62±0.19 folds, siSTX1A-2: 3.13±0.35 folds compared to siControl) (**Fig. 3I**). Interestingly, the size of lysosomes was increased in ionomycin treated control cells compared to its DMSO treatment (shControl-DMSO: 0.17±0.02 μm^2^ and shControl-ionomycin: 0.28±0.02 μm^2^) (**Fig. 3J**). However, the lysosome size was significantly decreased in STX1A knockdown compared to control cells upon ionomycin treatment (shControl: 0.28±0.02 μm^2^, shSTX1A-1: 0.21±0.01 μm^2^, and shSTX1A-2: 0.15±0.01 μm^2^) (**Fig. 3J**).

Next, we tested the lysosome exocytosis using LLOMe treatment or overexpression of lysosomal gene transcription factor, TFEB, in the cells (Zhu et al., 2020). As expected, treatment of STX1A knockdown cells with LLOMe showed enhanced lysosome dispersion above 15 μm (but not between 5-15 μm) distance, which is more than the methanol (control) treated cells (for 5-15 µm distance: siControl-methanol: 1.47±0.13, siSTX1A-methanol: 1.31±0.11, siControl-LLOMe: 2.30±0.21, siSTX1A-LLOMe: 2.00±0.21) (for above 15 µm: siControl-methanol: 0.35±0.03, siSTX1A-methanol: 0.53±0.06, siControl-LLOMe: 0.62±0.07, siSTX1A-LLOMe: 0.88±0.14) (**Fig. S4C** and **S4D**). Interestingly, TFEB-GFP localization was not altered with STX1A depletion compared to control cells (**Fig. S4E**). In line, the dispersion of lysosomes with TFEB-GFP overexpression in STX1A knockdown cells was not altered significantly compared to control cells (for 5-15 µm distance: siControl-non-transfected: 1.07±0.11, siSTX1A-non-transfected: 1.15±0.08, siControl-TFEB-GFP: 0.68±0.05, siSTX1A-TFEB-GFP: 0.75±0.10) (for above 15 µm: siControl-non-transfected: 0.26±0.03, siSTX1A-non-transfected: 0.51±0.06, siControl-TFEB-GFP: 0.18±0.02, siSTX1A-TFEB-GFP: 0.12±0.03) (**Fig. S4F**). Together, these results illustrate that the depletion of STX1A in HeLa cells leads to an inhibition in lysosome exocytosis, thereby stalling the lysosomes beneath the cell surface with an additional defect in protease activity.

### STX1A knockdown in HeLa cells decreases the degradative capacity of lysosomes and increases the number of autolysosomes

Next, we studied the status of late endosomes with respect to lysosomes upon STX1A depletion in HeLa cells. Cells were stained for late endosomal markers LBPA or CD63 along with LAMP1 and measured Pearson’s correlation coefficient between them. IFM analysis showed no change in the colocalization efficiency between LBPA and LAMP1 (**Fig. 4A** and **4B**) (siControl: *r*=0.52±0.03, siSTX1A-1: *r*=0.53±0.04, and siSTX1A-2: *r*=0.47±0.03). Similar results were obtained with CD63 and LAMP1 markers (**Fig. 4A**), suggesting that STX1A depletion does not affect the characteristics of late endosomes in HeLa cells. Interestingly, immunoblotting analysis showed no difference in the expression of several lysosomal proteins in STX1A knockdown compared to control cells (**Fig. 4C**), indicating no change in lysosome biogenesis in the cells. Correspondingly, the expression levels of lysosome transcription factors TFEB and TFE3 were also not affected dramatically in the cells (**Fig. 4C**). However, the cells displayed elevated levels of epidermal growth factor receptor (EGFR) and its phosphorylated form on immunoblots indicating a defect in EGFR degradation in STX1A knockdown compared to control cells (**Fig. 4D**). In contrast, the levels of transferrin receptor (TfR) showed no change in the cells (**Fig. 4D**). These studies suggest a defect in lysosome degradation of endocytosed cargo in the STX1A depleted compared to control HeLa cells.

Defective lysosome function alters the flux of autophagy, which generally accumulates the autophagosomes in the cells. Moreover, both lysosomes and autophagosomes localize closely at the perinuclear area for efficient fusion (Li Yu, 2018). We also hypothesized that STX1A knockdown cells display a slightly higher number of lysosomes in the cell periphery, which may result in reduced fusion of autophagosomes and thus cause the accumulation. To test this hypothesis, we expressed RFP-LC3 and found significantly increased number of autophagosomes in the STX1A depleted compared to control cells (**Fig. 4E** and **4F**) (number of puncta: shControl - 86±3, shSTX1A-1 - 111±6, and shSTX1A-2 - 113±5) (shSTX1A-1: 1.46±0.08 folds and shSTX1A-2: 1.40±0.05 folds compared to shControl). Interestingly, the Pearson’s correlation coefficient between RFP-LC3 and LAMP1 showed no significant change in the cells (siControl: *r*=0.33±0.02, siSTX1A-1: *r*=0.37±0.03, and siSTX1A-2: *r*=0.34±0.03) (**Fig. 4E** and **4F**). Consistently, the protein levels of LC3-II were not altered upon STX1A depletion compared to control cells (**Fig. 4G**). However, STX1A knockdown cells display enhanced expression of beclin, corresponding to an enhanced number of autophagosomes (**Fig. 4G**). These studies indicate an enhanced level of autophagy without affecting the fusion of autophagosomes with lysosomes. Towards this hypothesis, we overexpressed tandem tag autophagy reporter ptfLC3 in the cells and analyzed the autophagosomes and autolysosomes (**Fig. 4E**). IFM analysis showed a greater number of autolysosomes in STX1A knockdown compared to control cells (**Fig. 4E** and **4F**) (siControl: 16±4, siSTX1A-1: 63±10, and siSTX1A- 2: 72±17). These studies illustrate that STX1A depletion in HeLa cells may not affect autolysosome formation (Jahreiss et al., 2008) but results in reduced turnover of these organelles.

### SNAP23, SNAP25 and VAMP2 knockdown phenocopy the cellular phenotype of STX1A depleted HeLa cells

In neuronal cells, VAMP2 (R-SNARE) and SNAP25 (Qbc SNARE) act as cognate SNAREs of STX1A for the exocytosis of synaptic vesicles. During this fusion, STX1A has been shown to present at the PM and VAMP2 on secretory vesicles (Cupertino et al., 2016; Sudhof, 2014). However, our studies in HeLa cells showed a major cohort of STX1A on the lysosome membranes. We hypothesize that lysosomal STX1A could possibly be paired with one of the possible R-SNAREs (VAMP2, VAMP3, VAMP4, VAMP7, and VAMP8) of the PM. To identify the suitable cognate R- SNARE for lysosomal STX1A, we tested the relative position of the SNAREs by co-expressing GFP/mCherry-STX1A along with R-SNAREs (VAMP2, VAMP3, VAMP4, VAMP7 and VAMP8) tagged with pHuji/mCherry or GFP tagged at their N-terminus in HeLa cells (Chitirala et al., 2019). IFM analysis revealed that most of the R-SNAREs overlapped with STX1A, except GFP-VAMP2 and GFP-VAMP3 (**Fig. S5A**). However, we did not observe the PM localization of GFP-VAMP3 (Hu et al., 2007) compared to GFP-VAMP2 (**Fig. S5A**). To confirm our results, we overexpressed mCherry-STX1A^Δ25-146^ (N-terminal delete lacking regulatory domain which localizes majorly to the cell surface) along with GFP-VAMP2 and observed their colocalization at the cell surface compared to mCherry-STX1A and GFP- VAMP2 (**Fig. S5A** and **S5B**). Next, we tested the possible role of VAMP2 in regulating lysosome function by performing shRNA mediated knockdown in HeLa cells and observed the distribution of STX1A-positive lysosomes (**Fig. S5C**). As expected, the VAMP2 depleted cells showed enhanced lysosome distribution compared to shControl cells (**Fig. S5C**, quantified below). To identify the involvement of other SNAREs that may follow the pattern of STX1A or VAMP2 knockdown cells, we carried out shRNA mediated knockdown of selected SNAREs: SNAP23 and SNAP25 (Qbc); STX7 (Qa), STX8 (Qc), and VAMP7 (R) in HeLa cells and observed for a change in lysosome dispersion (**Fig. S5C**). Interestingly, we observed a change in lysosomal dispersion in cells depleted for SNAP23, SNAP25 and STX8. We proceeded with SNAP25 and VAMP2, as these SNAREs are known to form a complex with STX1A in neuronal cells (Cupertino et al., 2016). We also selected SNAP23 for our experiments as this SNARE shares homology with SNAP25 (Greaves et al., 2010) and displayed similar lysosome dispersion phenotype upon their gene knockdown separately in HeLa cells (**Fig. S5C**, quantified below).

From the above studies, we have selected VAMP2 (R), SNAP23 (Qbc) and SNAP25 (Qbc) genes along with STX1A (Qa) and validated their knockdown in HeLa cells (using lentiviruses) and subjected the cell lysates for immunoblotting analysis (**Fig. 5A** and **5B**). In addition to the reduced protein levels of individual SNAREs in their respective knockdown cells, we observed decreased STX1A stability in the VAMP2, SNAP23 and SNAP25 depleted condition (**Fig. 5B**). As expected, the VAMP2-, SNAP23- and SNAP25- knockdown cells showed the enhanced dispersion (measured by radial profile intensity of LAMP1) of lysosomes at a distance above 15 µm in reference to 0-5 µm distance, as similar to STX1A depleted cells (for 5-15 µm distance: shControl: 1.39±0.09, shSTX1A: 1.56±0.10, shVAMP2: 1.65±0.11, shSNAP23: 1.54±0.10, and shSNAP25: 1.64±0.14) (for above 15 µm: shControl: 0.31±0.03, shSTX1A: 0.54±0.06, shVAMP2: 0.56±0.06, shSNAP23: 0.50±0.04, and shSNAP25: 0.49±0.05) (**Fig. 5A** and **5C**).

**Figure 5.**
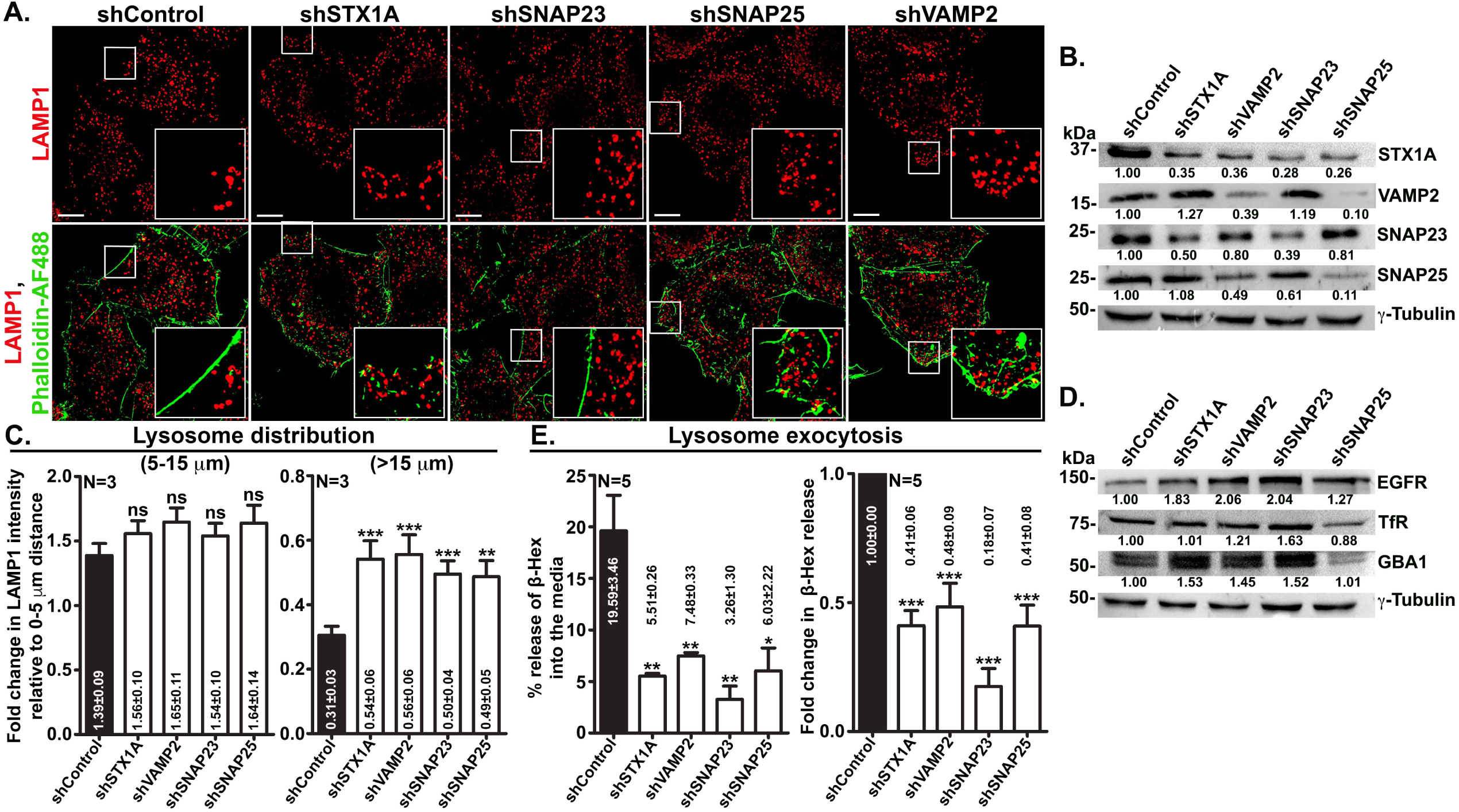
SNAP23, SNAP25 or VAMP2 knockdown phenocopy the STX1A depletion in HeLa cells. (A) IFM analysis of HeLa cells transduced with lentivirus containing shControl, shSTX1A, shSNAP23, shSNAP25 or shVAMP2. The cells were stained for LAMP1 and phalloidin-Alexa Fluor 488. Individual and merged panels are shown separately. Insets are magnified views of the white boxed areas. Scale bars, 10 µm. (B, D) Immunoblotting analysis of cell lysates. The efficiency of each gene knockdown is shown in B. The expression of different cargo in the knockdown cells is shown in E. γ-tubulin is used as the loading control in all immunoblots. The fold change in band intensities was normalized with internal control and indicated on the blots. (C) The plots represent the ratio of change in radial profile intensity of lysosomes between 5-15 µm or more than 15 µm distance with respective to 0-5 µm distance, respectively. (E) Plot represents the percentage change in the release of β-Hex into the supernatant (as a measure of lysosome exocytosis) of different SNAREs knockdown cells. Similarly, fold change in the release of β-Hex into the supernatant is shown separately. The average values in mean±s.e.m. are indicated on the graph. N=3 or 5. ns-non-significant, *p≤0.1, **p≤0.01 and ***p≤0.001.

Interestingly, an increase in EGFR but not transferrin receptor (TfR) levels was observed in the knockdown cell lysates of STX1A, VAMP2 and SNAP23. However, we observed a marginal increase in EGFR and reduced TfR protein levels in the SNAP25 knockdown HeLa cells (**Fig. 5D**). We also observed similar trend in the stability of lysosomal enzyme β-GC (GBA1) protein levels in the knockdown cell lysates (**Fig. 5D**). Next, we tested the lysosome exocytosis by performing the β-Hex activity assay in the supernatant of STX1A, VAMP2, SNAP23 and SNAP25 knockdown cells. As expected, there was a dramatic decrease in the percentage release of β-Hex activity in the supernatants upon knockdown of these SNAREs in HeLa cells (raw values: shControl: 19.59±3.46, shSTX1A: 5.51±0.26, shVAMP2: 7.48±0.33, shSNAP23: 3.26±1.30, and shSNAP25: 6.03±2.22) (in folds: shControl: 1.00±0.00, shSTX1A: 0.41±0.06, shVAMP2: 0.48±0.09, shSNAP23: 0.18±0.07, and shSNAP25: 0.41±0.08) (**Fig. 5E**). Moreover, subcellular fractionation revealed that STX1A, SNAP23, SNAP25, VAMP2 share the same member fractions (**Fig. S5D**), indicating a possibility of forming a SNARE complex between these SNAREs. Together, these results indicate that SNAP23 or SNAP25 and VAMP2 may act as STX1A cognate SNAREs in the lysosome exocytosis of HeLa cells.

### STX4A depletion in HeLa cells results in reduced STX1A levels and causes a moderate increase in lysosome dispersion

Previous studies have shown that STX4 and SNAP25 on PM pairs with VAMP7 on lysosomes during their exocytosis (Rao et al., 2004). We tested the role of PM localized STX4 in regulating lysosome function using siRNA-mediated gene knockdown in HeLa cells and validated it by semiquantitative RT-PCR analysis (**Fig. S6A**). Interestingly, we observed a dramatic loss of STX1A but not SNAP23/25 or VAMP2 transcripts in the STX4 knockdown cell lysate (**Fig. S6A**). In line with these results, both STX1A and SNAP23 protein levels (but not SNAP25 or VAMP2) were reduced in the STX4 depleted cells (**Fig. S6B**). Based on these results, we predicted that STX4 knockdown cells should mimic the phenotypes of STX1A depletion in HeLa cells. IFM analysis showed moderate dispersion of lysosomes beyond 15 μm distance with respect to 5-15 μm distance from the nucleus in STX4 depleted compared to control cells (for 5- 15 µm distance: shControl: 1.44±0.11 and shSTX4: 1.48±0.13; for above 15 µm: shControl: 0.41±0.04 and shSTX4: 0.58±0.04) (**Fig. S6C** and **S6D**). However, quantification of lysosome dynamics in STX4 knockdown cells showed a minor change in lysosome number (in fold change - shControl: 1.00±0.00 and shSTX4: 1.15±0.06) but not the lysosome size (shControl: 1.87±0.11 μm^2^ and shSTX4: 1.95±0.10 μm^2^) compared to control cells (**Figs. S6E** and **S6F**). In contrast to STX1A knockdown cells, STX4 depleted cells displayed no change in the protein levels of EGFR and GBA1 and enhanced LAMP1 protein levels compared to control cells (**Fig. S6B**). Overall, these studies suggest that STX4 depletion showed no effect on lysosome number, size and activity; however, the increased lysosome dispersion may be due to the loss of STX1A function in the cells.

### STX1A form SNAREpin complex with SNAP23/25 and VAMP2 in HeLa cells

To further probe the role of STX1A in lysosome exocytosis, we performed TIRF imaging microscopy of HeLa cells expressing LAMP1-GFP (label lysosomes). Interestingly, a large number of LAMP1-GFP vesicles are accumulated at the cell surface in STX1A depleted cells compared to control cells (**Fig. 6A**, **Videos S1** and **S2**). In line, quantification of the average number of LAMP1-GFP vesicles and their dynamic localization at the cell surface were significantly enhanced in STX1A knockdown compared to control cells (average number of spots: shControl: 17±1 and shSTX1A: 23±1; average number of spots/Sec/cell: shControl: 0.0041±0.0002 and shSTX1A: 0.0055±0.0002) (**Fig. 6A**, top graphs). Furthermore, we observed a significant decrease in the resident time of LAMP1-GFP vesicles in shSTX1A cells compared to shControl cells (average cell surface resident time (Sec)/cell: shControl: 3.92±0.22 and shSTX1A: 2.90±0.12; cell surface resident time in Sec (log_10_): shControl: 3.53±0.17 and shSTX1A: 2.74±0.11) (**Fig. 6A**, bottom graphs). These studies demonstrate that STX1A depletion in HeLa cells accumulates more LAMP1 vesicles at the cell surface with less resident time than in control cells. These studies will indirectly indicate a defect in the fusion of LAMP1 vesicles with the PM. Next, we tested the biochemical interaction of STX1A with its partner SNAREs in HeLa cells. As predicted, STX1A showed strong interaction with SNAP23, SNAP25 and VAMP2 but not with VAMP7, as well as with LAMP1 or EGFR (used as negative controls) (**Fig. 6B**). In conclusion, STX1A is possibly involved in the fusion of lysosomes (LAMP1- positive vesicles) with PM by paring with SNAP23 or SNAP25 and VAMP2. Future studies are required to investigate the specific conditions at which the individual SNARE SNAP23 or SNAP25 pairs with STX1A in mediating lysosome exocytosis.

**Figure 6.**
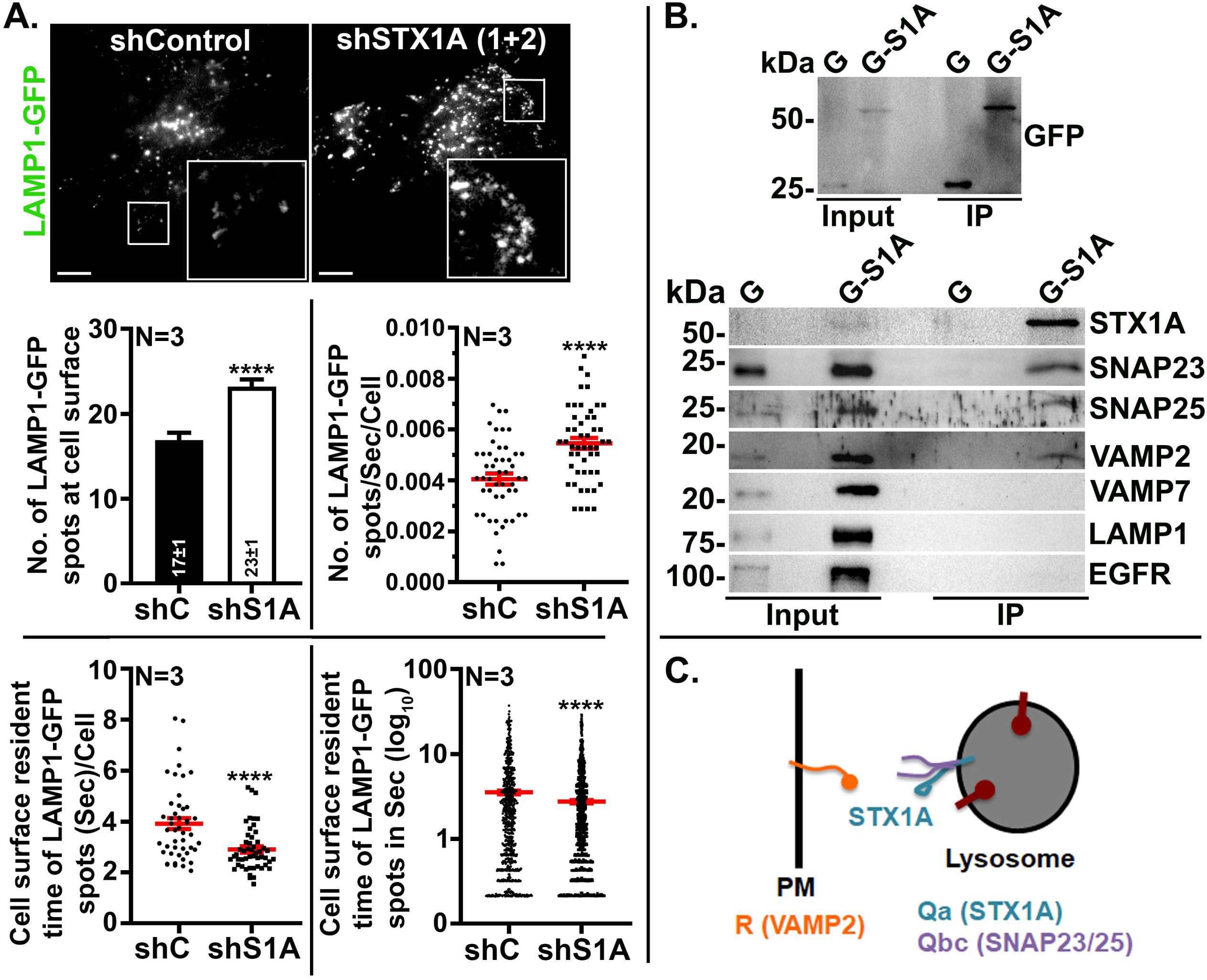
STX1A interacts with SNAP23, SNAP25 and VAMP2. (A) TIRF imaging of HeLa cells transduced with lentivirus containing shControl or shSTX1A mix (sh1+sh2). Cells were transfected with LAMP1-GFP and analyzed by live cell imaging. Insets are magnified views of the white boxed areas. Scale bars, 10 µm. Top: Plots represent the relative number of LAMP1- GFP vesicles at the cell surface and their kinetics in control and STX1A depleted cells. Bottom: Plots illustrate the resident time and the kinetics of LAMP1-GFP vesicles at the cell surface in control and STX1A knockdown cells. (B) GFP trap pulldown of GFP (G) or GFP-STX1A (G-S1A) expressing HeLa cell lysates. Pulldown samples were probed for input (top blot) and possible interacting SNAREs (bottom blot), including the negative controls such as VAMP7, LAMP1 and EGFR. IP – immunoprecipitated sample. (C) Model representing the lysosome exocytosis in HeLa cells. During this process, lysosome localized STX1A (Qa, blue) form a complex with SNAP23 or SNAP25 (Qbc, purple) and facilitate the fusion by interacting with plasma membrane (PM) SNARE VAMP2 (R, orange).

## Discussion

The Qa-SNARE STX1A in neuronal cells plays a critical role in releasing neurotransmitters through synaptic vesicle fusion with the PM. During this process, STX1A on the PM pairs with SNAP25 (Qbc) and interacts with synaptic vesicle SNARE VAMP2 (R) for the successful fusion (Cupertino et al., 2016; Jahn and Fasshauer, 2012; Sudhof, 2014). Even though the function and biochemical properties of STX1A have been studied extensively in the context of neurotransmitter release (Sudhof, 2014), its role in non-neuronal cells remains poorly understood. In this study, we attempted to comprehend the function of STX1A in HeLa cells. Fixed, live, and TIRF imaging and biochemical analyses demonstrated that STX1A uniquely localizes to the lysosomes and a cohort to the PM and late endosomes in HeLa cells. The depletion of STX1A caused the enhanced peripheral distribution and accumulated lysosomes just beneath the cell surface. Further, these cells exhibited a significant defect in lysosome exocytosis as measured by the β-Hex release assay and ionomycin induction assay. Additionally, STX1A depleted cells showed significantly decreased lysosome degradation with an enhanced number of autolysosomes. However, cargo delivery to the lysosomes was unaffected in STX1A knockdown cells. Interestingly, the N-terminal regulatory domain of STX1A plays an important role in its function and transport to the lysosomes. Moreover, we observed that STX1A partitioned to damaged lysosomes upon treatment with LLOMe, and its overexpression reduces the number of damaged lysosomes, suggesting a unique role for STX1A during lysosomal damage. Biochemically, STX1A pairs with SNAP23 or SNAP25 and VAMP2 and their respective depletion mimics the STX1 knockdown phenotypes in HeLa cells, suggesting that they might act as cognate SNAREs during lysosome exocytosis. These results placed STX1A as a crucial SNARE for the lysosomal exocytosis by localizing to these organelles.

The localization of STX1A to lysosomes in HeLa cells is unique to these organelles. Studies in MDCK cells reported an intracellular localization of STX1A, positive for lysosome marker AC17 but not to the PM (Low et al., 1996). Because STX1A expression has not been previously observed in MDCK cells, the authors concluded that the targeted STX1A may undergo lysosomal degradation (Low et al., 1996). In contrast, a proteomic analysis reported the expression of STX1A in HeLa cell lysate (Bekker-Jensen et al., 2017), and we confirmed it in this study via immunoblotting analysis (**Fig. S1A**). Further, our studies demonstrated the predominant localization of GFP-STX1A to lysosomes in HeLa cells and a cohort to the PM (**Figs. 1** and **S1**). The N-terminal regulatory domain of STX1A is important for its localization to the lysosomes, and its deletion mislocalizes and partly accumulates the SNARE in the Golgi (**Figs. 2** and **S2**). In line with our observations, a considerable cohort of N-terminal deleted STX1A mutant was observed with a Golgi marker in INS-1 cells (Yang et al., 2006). Additionally, a cohort of GFP-STX1A^Δ25-146^ was positive for lysosomes and the PM in HeLa cells (**Figs. 2** and **S2**). Interestingly, GFP-STX1A localization to lysosomes was reduced in Rab7 and Arl8b knockdown cells without affecting the STX1A protein levels (**Figs. 2** and **S2**). Consistently, the dominant negative mutant expression of Rab7 causes the clustering of STX1A around the perinuclear area, suggesting an indirect role of Rab7 in regulating the localization of STX1A to the lysosomes. Further, mCherry-STX1A localized more with the LLOMe-mediated damaged lysosomes marked by YFP-GAL3 compared to LAMP1-positive organelles and reduced their number significantly in the cells expressing mCherry-STX1A relative to non-expressing cells (**Fig. S2**). These studies indicated that Rab7, lysosomal damage and the N-terminus of STX1A possibly alter the STX1A-dependent lysosomal function by modulating its localization.

To understand the precise function of STX1A, we depleted STX1A in HeLa cells, which caused the dispersion of lysosomes, and they accumulated beneath the cell surface (marked by actin) relative to the control cells (**Figs. 3** and **S3**). However, STX1A knockdown cells did not display any difference in size or acidity but slightly enhanced lysosome number compared to control cells (**Fig. 3** and **S4**). The measurement of lysosomal enzyme activity (cathepsin B and DQ-Red BSA proteolytic activity assays) demonstrated a significant decrease in STX1A depleted compared to cells (**Figs. 3** and **S4**). Consistently, an enhanced accumulation of degradative cargo (EGFR) but not the recycling cargo (TfR) in addition to an increased number of autophagosomes in the STX1A depleted cells compared to control cells (**Fig. 4** and **S4**). In contrast, the lipid hydrolase activity (GBA activity) appears to be moderately increased in STX1A knockdown cells compared to control cells, and it may be due to increased protein levels or partly enhanced lysosomal number in the cells (**Figs. 3** and **4**). Interestingly, STX1A depleted cells showed no apparent defect in the fusion of autophagosomes or late endosomes with lysosomes; however, the increased number of autolysosomes possibly due to a difference in autophagic flux (**Fig. 4**). Overall, these results placed STX1A as an important component of lysosome function.

Recent studies highlighted the capability of releasing the lysosomal contents outside the cell through lysosomal exocytosis (Buratta et al., 2020; Tancini et al., 2020). This process involves the movement of lysosomes towards the cell periphery for the fusion with the PM and the release of lysosomal luminal content into the extracellular environment (Buratta et al., 2020). We tested these possibilities in the STX1A knockdown cells and observed that the cells disperse the lysosomes towards the periphery but decreased the release of lysosome content into the media (**Fig. 3** and **S4**). In line with these results, the percentage release of β-Hex into the cell media was drastically reduced upon induction of lysosome exocytosis with ionomycin in STX1A knockdown compared to control cells (**Fig. 3** and **S4**). As expected, the STX1A depleted cells showed increased lysosomes with enhanced peripheral distribution in ionomycin treated conditions (**Fig. 3** and **S4**). These studies demonstrate a possible role of STX1A in lysosome exocytosis, and its depletion blocks the fusion of lysosomes with PM by accumulating beneath the cell surface. Moreover, we predict that the N-terminal regulatory domain of STX1A may play a key role by an unknown mechanism during this process, and its deletion (GFP-STX1A^Δ25-146^) alters the lysosomal dynamics and follows similar phenotypes as the overexpression of GFP- STX1A in HeLa cells.

Complementary to the studies in neuronal cells (Cupertino et al., 2016; Sudhof, 2014), STX1A pairs with its cognate SNAREs SNAP25 (Qbc) or SNAP23 and VAMP2 (R) in HeLa cells for regulating lysosome exocytosis at basal level (**Fig. 6C**). In contrast to the reported study in NRK (normal rat kidney epithelial) cells, wherein VAMP7 (R) pairs with STX4 (Qa) and SNAP23 (Qbc) and form a *trans*-SNARE complex for the fusion of lysosome with PM (Jaiswal et al., 2002; Rao et al., 2004). In our hands, the knockdown of VAMP7 in HeLa cells did not show any lysosomal phenotypes (such as enhanced peripheral dispersion) compared to STX1A depletion in the cells (**Fig. S5**), relatively similar to the studies in B-lymphocytes (Obino et al., 2017). In line with this hypothesis, VAMP7 did not show any biochemical interaction with STX1A in HeLa cell lysates (**Fig. 6**). Further, our studies here (**Fig. S6**) clearly demonstrated that STX4 may not directly responsible for the lysosome exocytosis in HeLa cells in contrast to the previous report in NRK cells (Rao et al., 2004), wherein STX4 and SNAP25 on the PM pair with VAMP7 on lysosome in a TRPML1 Ca^2+^ release manner. We observed reduced STX1A levels in the STX4 depleted HeLa cells compared to control cells, and the siSTX4 cells follow the lysosome dispersion phenotype as similar to siSTX1A cells (**Figs. 3** and **S6**). However, STX4 knockdown cells did not show the accumulation of EGFR or GBA1 in contrast to shSTX1A cells (**Fig. S6**). Thus, we predict STX4 may not act as a Qa-SNARE during lysosome exocytosis in HeLa cells. However, the mechanism for reduced STX1A levels in STX4 depleted cells requires future investigation. Next, we analyzed the distinct role of SNAP25 and SNAP23 in lysosome exocytosis. However, we did not observe noticeable differences in cellular phenotypes between them or their interaction with STX1A (Figs. **5** and **6**). Moreover, SNAP23 showed around 60% identity with SNAP25 (Greaves et al., 2010). But SNAP25 and SNAP23 dock secretory granules to the PM at high (1 µm) and low (100 nm) Ca^2+^ levels, respectively, by interacting with different synaptotagmin family members (Chieregatti et al., 2004). Exogenous SNAP23 expression leads to high hormonal secretion under basal conditions in endocrine cells, suggesting that SNAP25 and SNAP23 expression levels possibly control the mode of granule release by forming docking complexes at different Ca^2+^ thresholds (Chieregatti et al., 2004; Sadoul et al., 1997). Hence, we predict that SNAP23 or SNAP25 acts as a putative SNARE partner of STX1A in lysosome exocytosis. Consistent with this hypothesis, the knockdown of SNAP25, SNAP23 and VAMP2 individually showed enhanced peripheral lysosome distribution and reduced β-Hex activity in the cell supernatants (**Fig. 5**). As similar to STX1A depleted cells, the accumulation of lysosomal degradative cargo EGFR was observed in the SNAP25, SNAP23 or VAMP2 knockdown cells (**Fig. 5**). However, the protein levels of GBA were increased in SNAP23 and VAMP2 knockdown cells but not in SNAP25 depleted cells in comparison to STX1A depletion, suggesting a possible preferable role of SNAP23 and VAMP2 in the lysosome exocytosis (**Fig. 5**). Moreover, GFP-VAMP2 localization was not observed with LAMP1-positive compartments (**Fig. S5**), providing a clue to act as R-SNARE during fusion by localizing to PM. Overall, these studies have provided a new set of SNAREs (**Fig. 6C**) that control the physiology of lysosomes during various cellular conditions.

## Materials and Methods

### Reagents and antibodies

All tissue culture reagents and chemicals were purchased from Sigma-Aldrich (Merck) or ThermoFisher Scientific (Invitrogen). 4-methylumbelliferyl β-D-glucopyranoside (MUD, GBA substrate, M3633), 4-methylumbelliferyl- N-acetyl-β-D-glucosaminide (β-hexosamindase substrate, M2133), ionomycin (I0634), LLOMe (L7393) and nocodazole (M1404) were procured from Sigma-Aldrich. Dextran-Alexa Fluor 647 (10K MW, D22914), dextran-Fluorescein (70K MW, D1822), DQ-Red BSA (D12051), lysoTracker Red DND-99 (L7528), phalloidin-Alexa Fluor 488 (A12379), phalloidin-Alexa Fluor 594 (A12381) and wheat germ agglutinin (WGA)- Alexa Fluor 594 (W11262) were obtained from ThermoFisher Scientific (Invitrogen). Matrigel was purchased from BD Biosciences.

The following commercial polyclonal and monoclonal antisera were used (m, mouse; h, human and r, rat proteins). Abcam: anti-LIMPII (ab16522), anti-Rab11 (ab128913), anti-Sodium Potassium ATPase (ab76020) and anti-VAMP7 (ab36195). BD Biosciences: anti-GM130 (610822) and anti-hp230/golgin-245 (611281). Cell Signalling Technology: anti-Rab9 (5118), anti-LAMP- 1 (9091), anti-EGFR (4267), anti-pEGFR (Tyr1068) (3777), anti-TFEB (4240s), anti-VAMP2 (13508) and anti-Beclin1 (3495). Cloud-Clone Corporation: anti-Cathepsin B (CTSB) (PAC964Mu01). Developmental Studies Hybridoma Bank: anti-LAMP-1 (H4A3), anti-LAMP-2 (H4B4), anti-CD63 (H5C6) and anti-EEA1 (3288). Invitrogen: anti-GFP (A-11122). ProteinTech: anti-Arl8b (13049-1-AP) and anti-Rab22A (12125-1-AP). Santa Cruz Biotechnology: anti-TfR (7087). Sigma-Aldrich: anti-Rab7 (R8779), anti-STX1A (SAB4502894), anti-Glucocerebrosidase (GBA1/GC) (G4171), anti-TFE3 (HPA023881), anti-γ- Tubulin (T6557), anti-LC3 (L7543) and anti-LBPA (MABT837). Synaptic systems: anti- SNAP23 (111202) and anti-SNAP25 (111002). All secondary antibodies were either from Invitrogen or Jackson Immunoresearch.

### siRNAs and plasmids

siRNAs: The following control siRNA (siControl) and target siRNA sequence against the respective gene were synthesized from Eurogentec, Belgium. siSTX1A-1: 5’- CCAGAAAGUUUGUGGAGGU-3’, siSTX1A-2: 5’-AGAACUCAUGUCCGACAUA-3’, siSTX1A-3’UTR: 5’-CACCAAAGGUCUUGGUACA-3’, siRab7: 5’- CUUCCCAUUUGUUGUGUUG-3’, siArl8b: 5’-GUCAAUGCUAUUGUUUACA-3’ and siSTX4: 5’-GGAGGAAGCUGAUGAGAACUAUAAC-3’.

shRNAs: The following shRNAs were from the human genome-wide TRC shRNA library, purchased from Sigma-Aldrich, USA (Catalog no. SH0411). pLKO.1-puro non-mammalian shRNA (referred to here as shControl, SHC002) was used as a control in all the shRNA mediated knockdown experiments. shSTX1A-1: 5’-AGAACTCATGTCCGACATAAA-3’, shSTX1A-2: 5’-CCAGAAAGTTTGTGGAGGTCA-3’, shSTX7-1: 5’-GCGATTATCAGTCTCATCATA-3’, shSTX7-2: 5’-CGTCTTATTCATGAGAGAGAA-3’, shSTX8-1: 5’- CCTCTATCATAAGTCGCCAAA-3’, shSTX8-2: 5’-CGTGACAATCAGAGCTTTGTT-3’, shSNAP23-1: 5’-GTGGATACATTAAACGCATAA-3’, shSNAP23-2: 5’- GAAACTCATTGACAGCTAAAG-3’, shSNAP25-1: 5’-CGATACACAGAATCGCCAGAT-3’, shSNAP25-2: 5’-GAACCATATCAACCAAGACAT-3’, shVAMP2-1: 5’- CATGAGGGTGAACGTGGACAA-3’, shVAMP2-2: 5’-CTGATTGAGGAAGGGTCTATT-3’, shVAMP4-1: 5’-GCATGTTCAGAATCAAGTGGA-3’, shVAMP4-2: 5’- GACCAAGATTTGGACCTAGAA-3’, shVAMP7-1: 5’-GCGAGTTCTCAAGTGTCTTAG-3’, and shVAMP7-2: 5’-TTGTATCACTGATGATGATTT-3’. A mix of two shRNAs (except shSTX1A) were used for the experiments.

Plasmids: pEGFP-C1-STX1A was constructed by cloning the PCR amplified STX1A from mouse cDNA prepared from mouse melanocytes (melan-Ink4a-Arf-1). The STX1A PCR product was digested with *Bgl*II and *Sal*I restriction enzymes and cloned into the same sites of the pEGFP-C1 vector. Similarly, pEGFP-C1-STX1A^Δ25-146^- was prepared by amplifying the truncated STX1A gene having 25-146 amino acids deletion followed by digestion with *Bgl*II and *Sal*I enzymes and then subcloned at the same restriction sites in pEGFP-C1 vector. The mCherry-C1-STX1A was made by subcloning the STX1A fragment from pEGFP-C1-STX1A by digesting with *Bgl*II and *Sal*I enzymes and then cloning into the same sites of the mCherry-C1 vector. The constructs were confirmed by restriction digestion and DNA sequencing.

Other plasmids: mCherry-UtrCh (26740), GFP-CD63 (62964), LAMP1-GFP (34831), LAMP1-RFP (1817), pMD2.G (VSV-G lentiviral envelop vector, 12259), psPAX2 (lentiviral packaging vector, 12260), pmRFP-LC3 (21075) and ptLC3 (rat LC3 fused to mRFP and EGFP, 21074) were obtained from Addgene. GFP-VAMP4, VAMP7-L-pHuji, VAMP8-mCherry, and GFP-VAMP3 were a kind gift from Jens Rettig, Saarland University, Homburg, Germany (Chitirala et al., 2019). Arl8b-Tomato, GFP-Rab7b^WT^, GFP-Rab7b^Q67L^ and GFP-Rab7b^T22N^ mutants were a kind gift from Mahak Sharma, IISER Mohali, India (Khatter et al., 2015). TFEB-GFP (Mahanty et al., 2019) was described previously. GFP-VAMP2 was a kind gift from Andrew A. Paden, The University of Sheffield, Sheffield, UK. YFP-GAL3 was a kind gift from Santosh Chauhan, CCMB, Hyderabad, India.

### Tissue culture, transfection and lentivirus viral transduction

The cell lines HeLa and HEK293T were (from ATCC) cultured in complete DMEM medium, consisting of DMEM (Invitrogen), 10% FBS (Biowest), 1% Pen-Strep (Invitrogen), and 1% L-glutamine (Invitrogen). The cells were maintained in a humidified cell culture incubator at 37°C with 10% CO_2_. The transfection of plasmids into the mammalian cells was carried out by starving the cells for 30 min in OPTI-MEM (Invitrogen) medium followed by adding the transfection mix having DNA (1 µg) and Lipofectamine 2000 (3µl; Invitrogen) to the cells in a 35 mm dish. After 6 hours, the media was replaced with a complete DMEM medium and incubated for 48 h at 37°C before use. Cells were fixed, immunostained, and imaged. For siRNA-mediated knockdown studies, HeLa cells were seeded at 60-70% confluency and incubated with a premix of 100 nM siRNA and 5 µl of oligofectamine (Invitrogen) in the OPTI-MEM medium. The cells were then incubated in complete DMEM at 37°C for a total of 48-52 hours before use. For IFM experiments, cells were plated on coverslips post 24 h of transfection, fixed, stained and then imaged. The preparation of lentivirus encoding shRNA and their infection of mammalian cells has been described previously (Shakya et al., 2018). In some experiments, cells were treated with 5 μM of nocodazole for 90 min or 1 mM LLOMe or methanol for 1 h at 37°C in a CO_2_ incubator. For ionomycin treatment experiments, cells were incubated in HBSS containing 10 μM ionomycin with 4 mM CaCl_2_ for 10 min in a CO_2_ incubator at 37°C.

Rescue of STX1A knockdown in HeLa cells was achieved by transfecting 3’-UTR specific siRNA followed by transfecting the cells with GFP-STX1A and incubated for 24 h. Post transfection, cells were stained and subjected to IFM or immunoblotting analyses.

### Internalization of dextran beads

HeLa cells were incubated with 0.5 mg/ml of dextran-fluorescein (70,000 MW) or dextran-Alexa Fluor 647 (10,000 MW) beads in a complete medium on ice for 30 min. The cells were then incubated for an additional 2 h at 37°C in a CO_2_ incubator. Finally, cells were washed with 1X PBS, fixed with 3% formaldehyde, immunostained and then imaged.

### Cell surface binding of wheat germ agglutinin-Alexa Fluor 594 (WGA-AF594)

The cells on coverslips were subjected to a wash and subsequently incubated with WGA-AF594 (0.5 ng/µl) in 200 µl of ice-cold HBSS buffer. The incubation was carried out for 15 min on ice under dark conditions. After incubation, the cells were washed with 1X PBS and fixed with 3% formaldehyde for 15 min at room temperature. Further, the cells were washed, stained and then analyzed using IFM.

### Immunofluorescence microscopy (IFM)

The cells seeded on coverslips were fixed with 3% formaldehyde for 15 - 20 min at room temperature. Subsequently, the coverslips were washed and preserved in 1X PBS containing 0.05% sodium azide until required. The cells were then incubated with primary antibodies and washed with 1X PBS, followed by staining with their corresponding secondary antibodies, as described previously (Jani et al., 2015). The colocalization efficiency (Pearson’s correlation coefficient, *r*) between the two proteins was calculated using the cellSens Dimension software (Olympus). For other experiments, cells seeded on coverslips were subjected to the internalization of either DQ-BSA-Red or dextran-fluorescein/Alexa Fluor 647. This was followed by the fixation of cells with 3% formaldehyde. In other experiments, cells were stained with either phalloidin-Alexa Fluor 488 or 594 (2 U/ml) in 1XPBS or wheat germ agglutinin (WGA)-Alexa Fluor 594 (50 μg/ml) in HBSS for a duration of 30-60 min. Subsequently, the cells were analyzed using IFM, using the microscope setup previously described (Jani et al., 2015).

For live cell imaging, cells were seeded in live cell dishes (MatTek Life Sciences) and then transfected with respective constructs as described. Cells were imaged under the Olympus IX81 wide-field microscope, and the movie was captured for one to two minutes.

### TIRF live cell imaging microscopy

The cells seeded on glass microwell live cell dishes (MatTek Life Sciences) were transfected with LAMP1-GFP. Post 24h, cells were imaged under a Nikon Ti2 spinning disk inverted fluorescence microscope using TIRF mode with a 100X objective at 61.5° critical angle (Tc). LAMP1-GFP puncta at the cell surface was captured in a movie format. Quantitative analysis (the number and residence time of LAMP1-GFP vesicles) of the acquired videos was performed using the TrackMate v.7.7.2 plugin within Fiji software. An estimated object diameter of 0.5 microns was applied by setting the quality threshold to 90%. The initial spot detection thresholding was adjusted to identify approximately 500 spots per cell. Each detected spot was tracked over time, and the resulting trajectories were plotted for further analysis.

### LysoTracker Red uptake and DQ-Red BSA trafficking assays

The cells on a coverslip were subjected to incubation for 1 h in a medium containing 100 nM of LysoTracker Red at 37°C. Subsequently, the cells were fixed using 4% paraformaldehyde. Similarly, the cells were incubated in a medium containing 10 μg/ml of DQ-Red BSA for 12 h at 37°C. The cells were further chased in a plain medium for 6 h before the fixation with 3% paraformaldehyde. The cells were then stained with anti-LAMP-1 antibody and then imaged subsequently.

### Quantification methods of IFM

*1. Quantification of the distribution of lysosomes:* The Radial Intensity Profile plugin was used in the FIJI software to calculate the particle distribution. The cell was divided into three distinct areas, including 0-5 μm, 5-15 μm and greater than 15 μm proximity from the nucleus. The radial intensity of these areas was then determined. The intensity profile of the areas 5-15 μm and greater than 15 μm were normalized with the intensity of the 0-5 μm area for each cell. These values were plotted as a measure of fold change in radial profile intensity to 0-5 μm area.
*2. Quantification of fluorescence intensity by corrected total cell fluorescence (CTCF):* The maximum intensity projection (MIP) of the image was generated by merging the layers using ImageJ. For this, select the Image tab>> select the stacks>> select the z-projection >> set the range of z-stacks >> click OK. Next, measure the area of the cell, mean fluorescence, and the integrated density. For this, go to Analyze>>set measurement >> select Area, Mean Grey Value and Integrated Density. Next, using the freehand tool, select the individual cell in the field and take note of the parameters. Further, go to Analyze>> Measure, which generates the data in a separate sheet that can be copied to an Excel file. The background intensity was calculated for each quantified image using the same method. The CTCF value for each cell was calculated using the formula: Integrated Density - (Area of the cell x Mean Fluorescence of the background). The average CTCF for each set was calculated and then plotted.
3. *Quantification of the number and size of fluorescent puncta:* First, open the images in ImageJ and then select the single channel (image tab>> color dropdown>> split channel). Next, merge the z-stacks into a single plane (image tab>> stacks dropdown>> z-projection). Convert the selected single-channel MIP image to 8 bit. Further, use the auto thresholding (in the Image tab) tool and adjust the image with a default range of 30 to 255. Then, convert the image to a binary using the Make Binary tool (in the Process tab). Perform watershed segmentation with the Watershed tool in the Process tab. Use the freehand selection tool in the Analyze tab to select the cell area and then analyze particles. The obtained puncta counts are based on default values of particle size and circularity. Calculate the average puncta number per cell and then plotted. Similarly, the average size of the puncta was calculated and then plotted. Additionally, the Fiji plugin ComDet can be used in certain cases to calculate the number of puncta (Patel et al., 2023a).
4. *Quantification of autophagosomes and autolysosomes:* Images of cells expressing ptfLC3 were analyzed. Yellow puncta represent the autophagosomes (GFP-RFP positive) and RFP-positive puncta represent the autolysosomes. The number of RFP- and GFP- positive puncta were quantified using the ComDet plugin in Fiji software. The number of autolysosomes was quantified by counting the RFP puncta alone. The number of autophagosomes was determined by subtracting RFP-positive puncta from the total number of GFP-positive puncta count. Both data were plotted separately.

### β-glucocerebrosidase (GBA) activity and cell viability assays

Cells were plated in triplicate wells of a 96 well clear flat-bottom black plate (Corning) and used for assays upon reaching a 70-80% confluency as described previously (Patel et al., 2023b). For cell viability assay, the cells were incubated in a growth medium containing 1 mg/ml of resazurin at 37°C for 3-4 hours. The fluorescence intensity (excitation at 530 nm and emission at 590 nm) was measured using a Tecan multi-mode plate reader (Infinite F200 Pro). After the resazurin assay, the cells were washed with 1X PBS and incubated in 50 μl of 3 mM MUD (4-methylumbelliferyl-β-D-glucopyranoside, made in 0.2 M sodium acetate buffer pH 4.2) at 37°C. The assay was stopped with 150 μl of 0.2 M glycine buffer pH 10.8 after 150 min (2.5 h) of incubation. The liberated 4-methylumbelliferone fluorescence intensity (excitation at 365 nm and emission at 445 nm) was measured using a Tecan multi-mode plate reader. The lysosomal enzyme activity (in A.U.) per well was then normalized with respective cell viability values. The fold change in GBA activity was calculated by comparing the values obtained with STX1A knockdown cells to the control cells and then plotted. The assay was performed trice before plotting the data.

### β-hexosaminidase (β-Hex) release (lysosome exocytosis) assay

We followed the protocol described previously in (Escrevente et al., 2021). Cells were incubated in HBSS containing 10 µM ionomycin and 4 mM CaCl_2_. After the incubation, cells were placed on ice and collected the supernatants, and then lysed the cells with 1% IGEPAL (prepared in dH2O at a 1:5 ratio) solution. The cell supernatants and lysates were incubated with 4-methylumbelliferyl-N-acetyl-β-D-glucosaminide (4-MU-β-D-GlcNAc) for 15 min at 37°C. Fluorescence intensity was measured using an Infinite F200 Pro plate reader (excitation at 365 nm and emission at 450 nm). Simultaneously, the protein content from the cell supernatants and lysates was determined using the BCA protein assay kit. HBSS and diluted 1% IGEPAL (1:5) were used as negative controls in the assay. The β-hexosaminidase (β-hex) activity was calculated for each sample by normalizing the activity with the total protein amount. β-hex activity in supernatant=(fluorescence (365/450)−HBSS alone)/protein (in µg). β-hex activity in cell lysate=(fluorescence (365/450)−IGEPAL alone)/protein (µg). Total β-hex activity=β-hex activity in supernatant+5× β-hex activity in cell lysate. Finally, the percentage of β-hex release was calculated as follows: β-hex release (% of total)=100×(β-hex activity in the supernatant/total β-hex activity).

### Isolation of RNA and preparation of cDNA

Total RNA from HeLa cells was isolated using a GeneJET RNA purification kit (ThermoFisher Scientific) in the presence of β-mercaptoethanol (Shakya et al., 2018). The RNA concentration was estimated using a NanoDrop 2000C spectrophotometer (ThermoFisher Scientific). The cDNA was prepared from total RNA using the RevertAid First Strand cDNA synthesis kit (K1622, ThermoFisher Scientific). Transcript analysis was performed as described previously (Jani et al., 2015).

### Immunoblotting

The cell pellets were lysed in SDS buffer and subjected to immunoblotting as described previously (Shakya et al., 2018). Protein concentrations were measured using Bradford reagent (Bio-Rad) and subjected to SDS-PAGE gel electrophoresis by loading equal amounts of lysates. The immunoblots were developed with Clarity Western ECL substrate (Bio-Rad) and captured images on a Molecular Imager ChemiDoc XRS+ imaging system (Bio-Rad). Subsequently, we analyzed the blots using Image Lab 4.1 software. Protein band intensities were quantified using the Image Lab software and normalized with γ-tubulin. The protein band intensities were represented as fold change relative to their respective control.

### Subcellular membrane fractionation

The fractionation was carried out by following the protocol described previously (Shakya et al., 2018). Briefly, HeLa cells were harvested, pelleted, washed once with ice-cold PBS, and homogenized in a lysis buffer (10 mM HEPES pH 7.4, 0.25 M sucrose, 1 mM EDTA, and 1× protease inhibitor cocktail) at 4°C using a Dounce homogenizer. The lysates were clarified by centrifugation at 800 *g* for 10 min at 4°C. Subsequently, the homogenates were fractionated on a manually layered sucrose step gradient (1.0, 1.2, 1.4, 1.6, 1.8, and 2.0 M sucrose) by ultracentrifugation at 120,000 *g* for 1 h at 4°C in an SW60Ti rotor using Beckman Coulter optima XPN-100 ultracentrifuge. Two fractions were collected per each step gradient from top to bottom and analyzed by immunoblotting.

### GFP-TRAP of STX1A pull down

The cell lysate for immunoprecipitation was prepared by lysing the cells in ice-cold lysis buffer (10 mM Tris-Cl pH 7.5, 150 mM NaCl, 0.5 mM EDTA, 0.5 % IGEPAL, adjust the pH at 4°C), supplemented with protease inhibitors to prevent the protein degradation. After 30 min of incubation on ice with occasional pipetting, the lysate was centrifuged and collected the supernatant. Beads were equilibrated in a dilution buffer (10 mM Tris-Cl pH 7.5, 150 mM NaCl, 0.5 mM EDTA, adjust the pH at 4°C), followed by an overnight incubation with the lysate at 4°C. Post binding, the beads were washed multiple times with wash buffer (10 mM Tris-Cl pH 7.5, 150 mM NaCl, 0.05 % IGEPAL, 0.5 mM EDTA, adjust the pH at 4°C) and eluted the bound proteins with 2X SDS-sample buffer, subjected to SDS-PAGE followed by immunoblotting.

### Statistical analyses

Statistical analyses were performed using GraphPad Prism 6.04. All experiments were conducted in triplicate, and the number of cells is presented in the graphs. Significance between groups was assessed using an unpaired Student’s t-test, with *P≤0.05, **P≤0.01, ***P≤0.001, ****P≤0.0001, and ns indicating non-significance.

## Supporting information

The live-cell TIRF imaging of siControl HeLa cells expressing LAMP1-GFP. Cells were imaged for approximately 2 min.

The live-cell TIRF imaging of siSTX1A HeLa cells expressing LAMP1-GFP. Cells were imaged for approximately 2 min.

## Acknowledgements

We thank Bishal Singh for his experimental support. We thank Praneeth Chitirala and Jens Retting (Saarland University, Homburg, Germany) for the generous gift of SNARE constructs. This work was supported by the Department of Biotechnology (BT/PR32489/BRB/10/1786/2019), Science and Engineering Research Board (CRG/2019/000281), DBT-NBACD (BT/HRD-NBA-NWB/38/2019-20), India Alliance (500122/Z/09/Z), and IISc-DBT partnership programme. Infrastructure in the department was supported by DST-FIST, DBT, and UGC. AMB was supported by DBT-JRF (DBT/2015/IISc/NJ-02).

## Author contributions

AMB performed all the experiments of this study. SRGS designed, oversaw the entire project, coordinated and discussed the work with the author.

## Conflict of interest

The authors declare that they have no conflict of interest.

## Supplementary information

### Supplementary Figures

**Figure S1.**
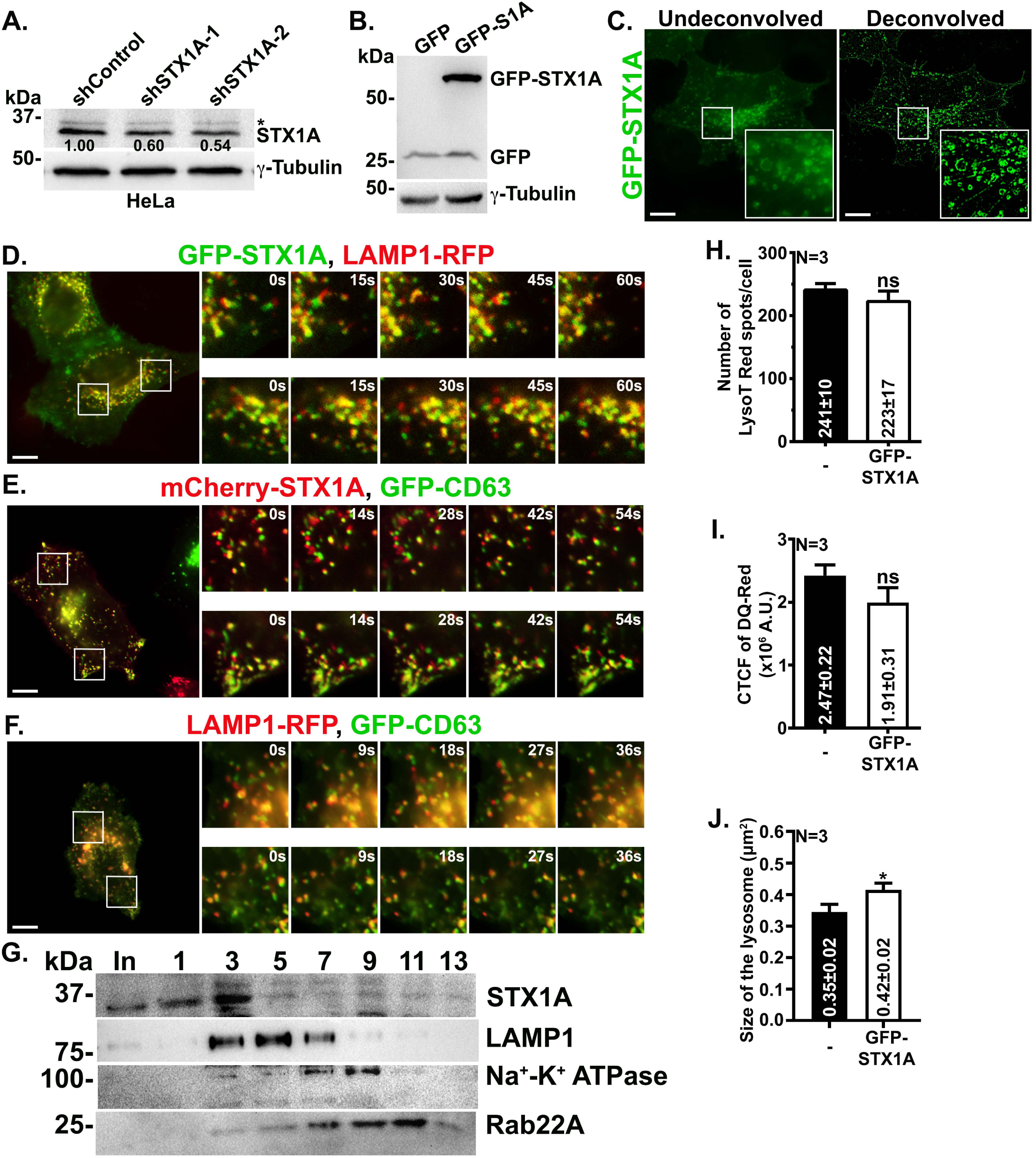
(A) Immunoblotting analysis of STX1A knockdown and control cells for checking the expression of STX1A. * non-specific band highlighted by anti-STX1A antibody. The fold change in band intensities was normalized with internal control and indicated on the blots. (B) Immunoblotting analysis of GFP-STX1A overexpression in HeLa cells. GFP expression is used as a control. The blot was probed with an anti-GFP antibody. γ-tubulin is used as the loading control in all immunoblots. (C) Images represent the GFP-STX1A overexpression in HeLa cells. Undeconvolved (left) and deconvolved (right) images are shown separately. Insets are magnified views of the white boxed areas. (D-F) Live cell imaging microscopy of HeLa cells expressing GFP-STX1A and LAMP1-RFP in D; mCherry-STX1A and GFP-CD63 in E; and GFP-CD63 and LAMP1-RFP in F. Insets are magnified views of the white boxed areas shown in the images at different time points (s, Sec). Scale bars, 10 µm. (G) Subcellular membrane fractionation of HeLa cells. Fractions were probed for STX1A, LAMP1 (lysosome marker), Na^+^-K^+^ ATPase (plasma membrane marker) and Rab22A (recycling endosome marker). In- Input and numbers represent the membrane (alternative) fractions from top to bottom of the sucrose gradient. (H) Plot represents the number of LysoTracker puncta in non-transfected (-) and GFP-STX1A overexpressing cells. (I) Plot represents the CTCF (A.U.) of DQ-Red intensity in non-transfected (-) and GFP-STX1A overexpressing cells. (J) Plot represents the size of lysosomes (μm^2^) in non-transfected (-) and GFP-STX1A overexpressing cells. The average values in mean±s.e.m. are indicated on the graphs. N=3. ns-non-significant and *p≤0.1.

**Figure S2.**
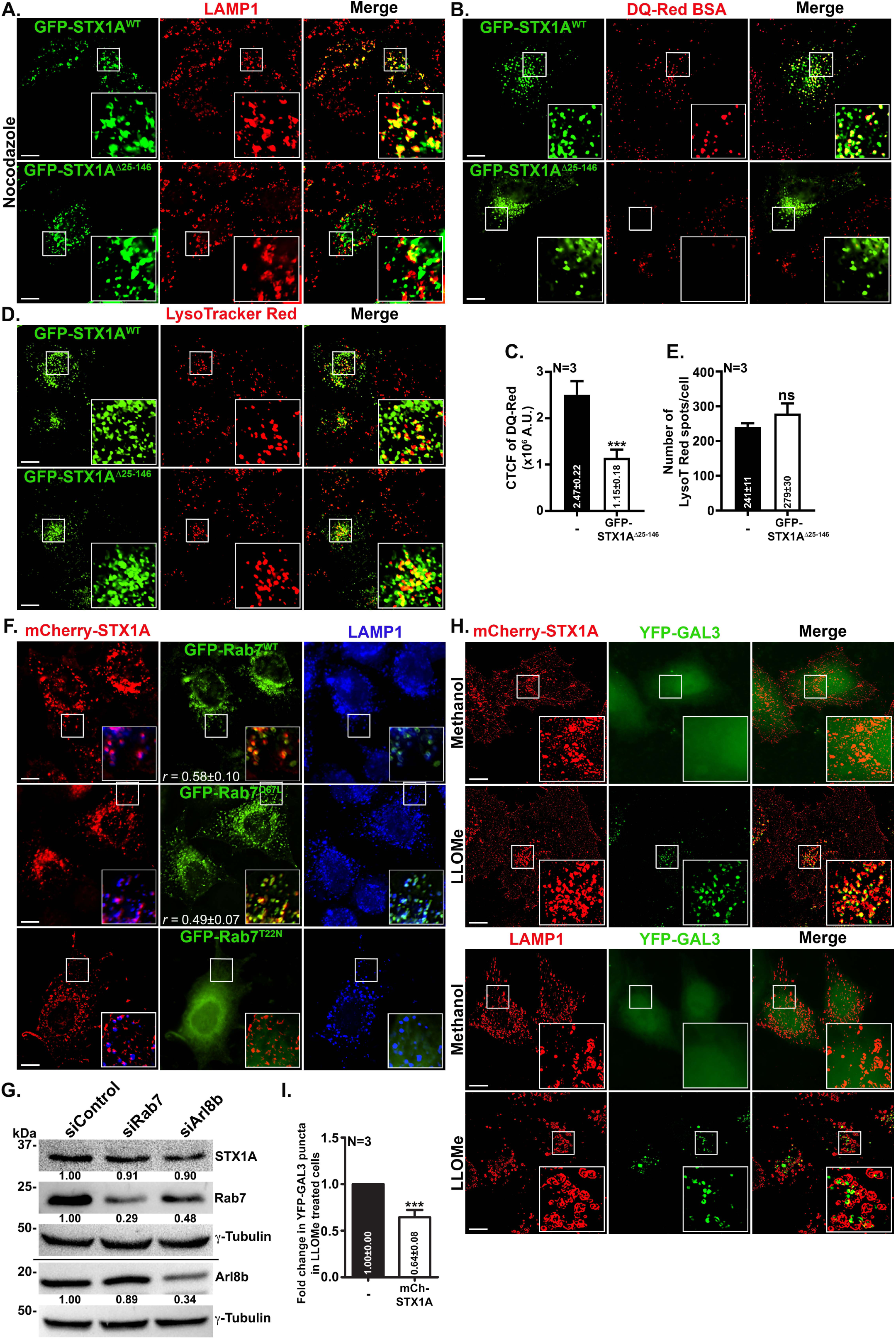
(A, B, D) IFM analysis of HeLa cells expressing GFP-STX1A^WT^ or GFP-STX1A^Δ25-146^. Cells were treated with nocodazole for 1.5 h, fixed and stained with anti-LAMP1 antibody in A. Cells were subjected to DQ-Red BSA processivity assay in B. Cells were subjected to LysoTracker-Red uptake assay in D. (C) Plot represents the CTCF (A.U.) of DQ-Red intensity in non-transfected (-) and GFP-STX1A^Δ25-146^ expressing cells. (E) Plot represents the number of LysoTracker-Red spots/cell in non-transfected (-) and GFP-STX1A^Δ25-146^ expressing cells. (F) IFM analysis of HeLa cells co-expressing mCherry-STX1A with GFP-Rab7^WT^, GFP-Rab7^Q67L^, or GFP-Rab7^T22N^. Cells were fixed and stained for LAMP1 (blue). The colocalization efficiency (*r*) between Rab7 and STX1A is indicated on the images. (G) Immunoblotting analysis of Rab7 and Arl8b knockdown cell lysates along with control cell lysate. Cell lysates were probed for the expression of STX1A. γ-tubulin is used as a loading control. (H) IFM analysis of HeLa cells transfected with YFP-GAL3 alone or along with mCherry-STX1A. After 24 h, cells were treated with either methanol or 3 mM LLOMe for 4 h followed by fixation. YFP-GAL3 transfected cells were stained for LAMP1. Insets are magnified views of the white boxed areas. Scale bars, 10 µm. (I) Plot represents the fold change in the number of YFP-GAL3 particles in non-transfected (-) and mCherry-STX1A expressing cells. Cells were treated with LLOMe. The average values in mean±s.e.m. are indicated on the graphs. N=3. ns-non-significant and ***p≤0.001.

**Figure S3.**
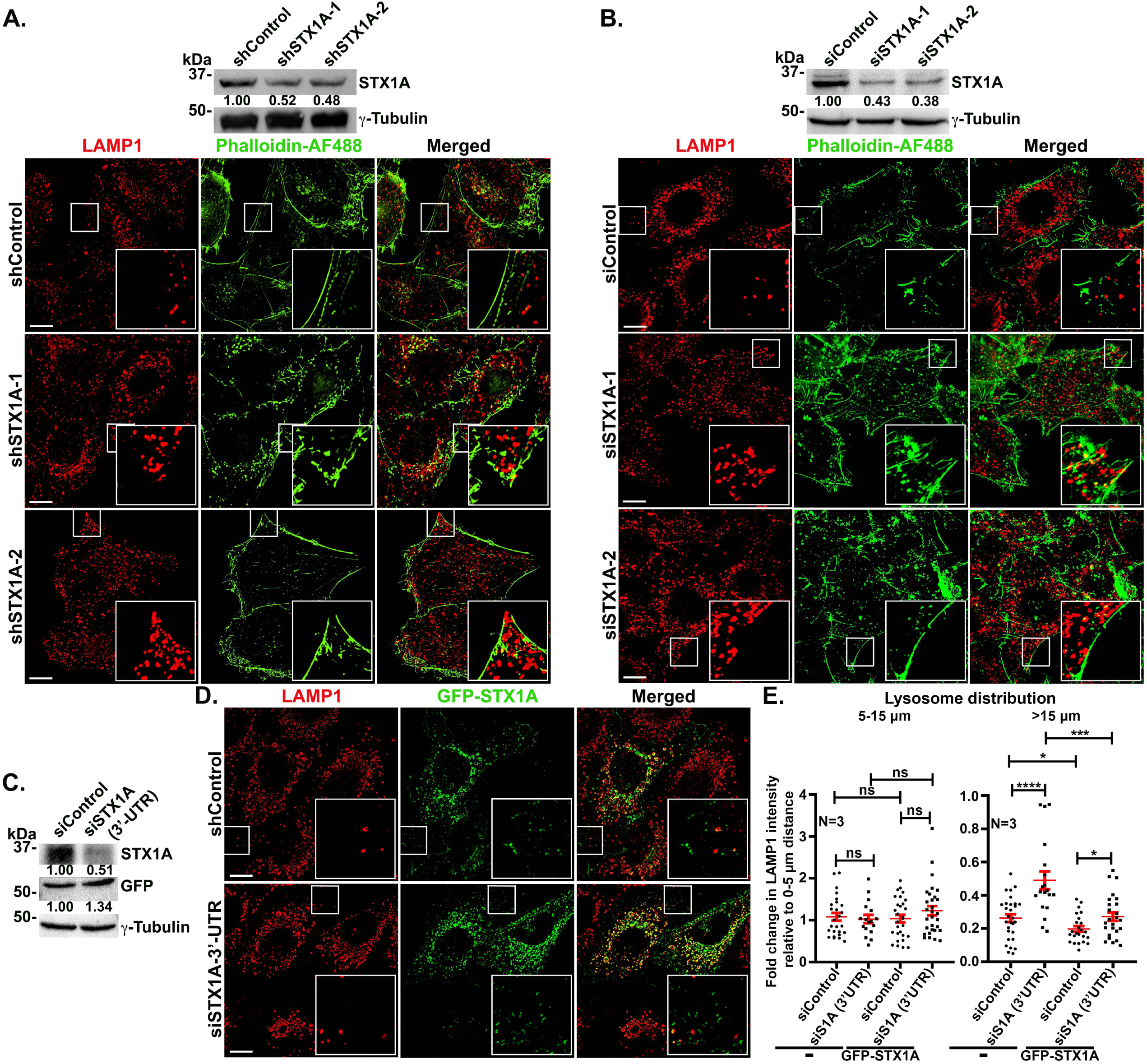
(A) Immunoblotting analysis of HeLa cell lysates from shControl, shSTX1A-1 and shSTX1A-2 cells. The blot was probed for the expression of STX1A. IFM analysis HeLa cells transduced with lentivirus containing shControl, shSTX1A-1, and shSTX1A-2. Cells were stained with anti-LAMP1 antibody and phalloidin-Alexa Fluor 488, followed by IFM. (B) Immunoblotting analysis of HeLa cell lysates from siControl, siSTX1A-1 and siSTX1A-2 cells. The blot was probed for the expression of STX1A. IFM analysis of HeLa cells transfected with siControl, siSTX1A-1 and siSTX1A-2, fixed and stained with anti-LAMP1 antibody and phalloidin-Alexa Fluor 488. (C) Immunoblotting analysis of siControl and siSTX1A (3’-UTR) HeLa cell lysates expressing GFP-STX1A. Blots were probed with anti-STX1A and anti-GFP antibodies separately. γ-tubulin is used as a loading control in all immunoblotting experiments. (D) IFM analysis of HeLa cells transfected with control or STX1A (3’-UTR) siRNAs. Cells were transfected with GFP-STX1A after 24 h. Insets are magnified views of the white boxed areas. Scale bars, 10 µm. (E) Plots represent the ratio of change in radial profile intensity of lysosomes between 5-15 µm or more than 15 µm distance with respect to 0-5 µm distance, respectively, as a measure of lysosome dispersion in the given conditions. -, non-transfected cells. N=3. ns-non-significant, *p≤0.1, ***p≤0.001 and ****p≤0.0001.

**Figure S4.**
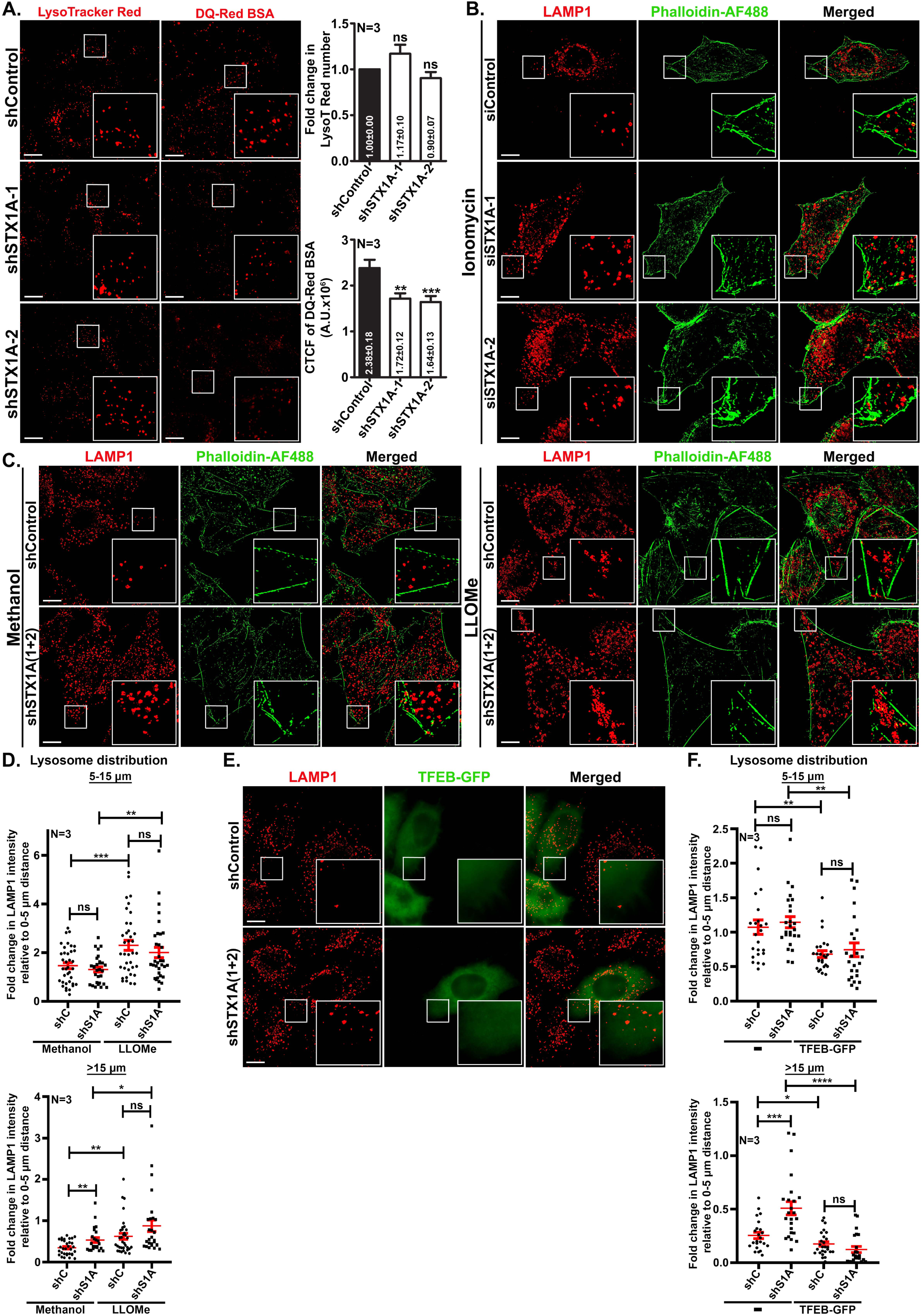
(A) IFM analysis of HeLa cells transduced with lentivirus containing shControl, shSTX1A-1 or shSTX1A-2 plasmids. Cells were subjected to the uptake of LysoTracker Red (left panel) or DQ-Red BSA (right panel), fixed and then imaged. Top plot represents the fold change in the number of LysoTracker-Red puncta/cell in STX1A knockdown compared to control cells. Bottom plot represents the CTCF (A.U.) of DQ-Red intensity in the control and STX1A depleted cells. The average values in mean±s.e.m. are indicated on the graphs. (B) IFM analysis of STX1A knockdown and control cells that were treated with ionomycin, fixed and stained with anti-LAMP1 antibody and phalloidin-Alexa Fluor 488. (C) IFM analysis of STX1A knockdown and control cells that were treated with methanol or LLOMe, fixed and stained with anti-LAMP1 antibody and phalloidin-Alexa Fluor 488. (D) Plots represent the ratio of change in radial profile intensity of lysosomes between 5-15 µm or more than 15 µm distance with respect to 0-5 µm distance, respectively, as a measure of lysosome dispersion in the given conditions (as shown in C). (E) IFM analysis of TFEB-GFP transfected STX1A knockdown and control cells, fixed and staining with anti-LAMP1 antibody. In all IFM images, insets are magnified views of the white boxed areas. Scale bars, 10 µm. (F) Plots represent the ratio of change in radial profile intensity of lysosomes between 5-15 µm or more than 15 µm distance with respect to 0-5 µm distance, respectively, in the given conditions (as shown in E). -, non-transfected cells. N=3. ns-non-significant, *p≤0.1, **p≤0.01, ***p≤0.001 and ****p≤0.0001

**Figure S5.**
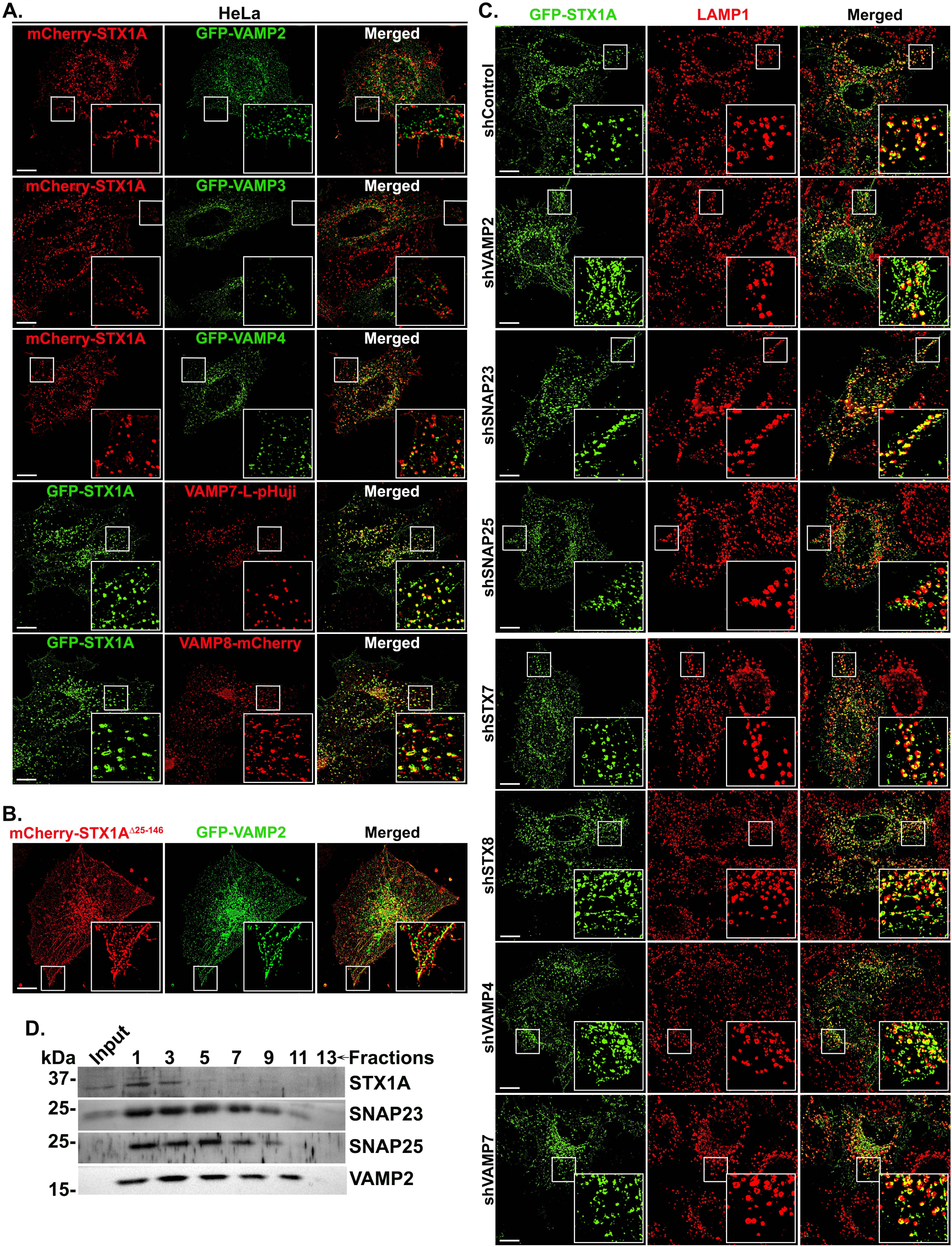
(A) IFM analysis HeLa cells co-expressing GFP-STX1A or mCherry-STX1A with GFP-VAMP2, GFP-VAMP3, GFP-VAMP4, VAMP7-L-pHuji or VAMP8-mCherry. (B) IFM analysis of HeLa cells transfected with mCherry-STX1A^Δ25-146^ and GFP-VAMP2. (C) IFM analysis HeLa cells transduced with lentivirus containing shControl, shVAMP2, shSNAP23, shSNAP25, shSTX7, shSTX8, shVAMP4 or shVAMP7 plasmids. The cells were overexpressed with GFP-STX1A and stained with anti-LAMP1 antibody. In all IFM images, insets are magnified views of the white boxed areas. Scale bars, 10 µm. (D) Subcellular membrane fractionation of HeLa cells. Fractions were probed for STX1A, SNAP23, SNAP25 and VAMP2. The numbers represent the membrane (alternative) fractions from top to bottom of the sucrose gradient.

**Figure S6.**
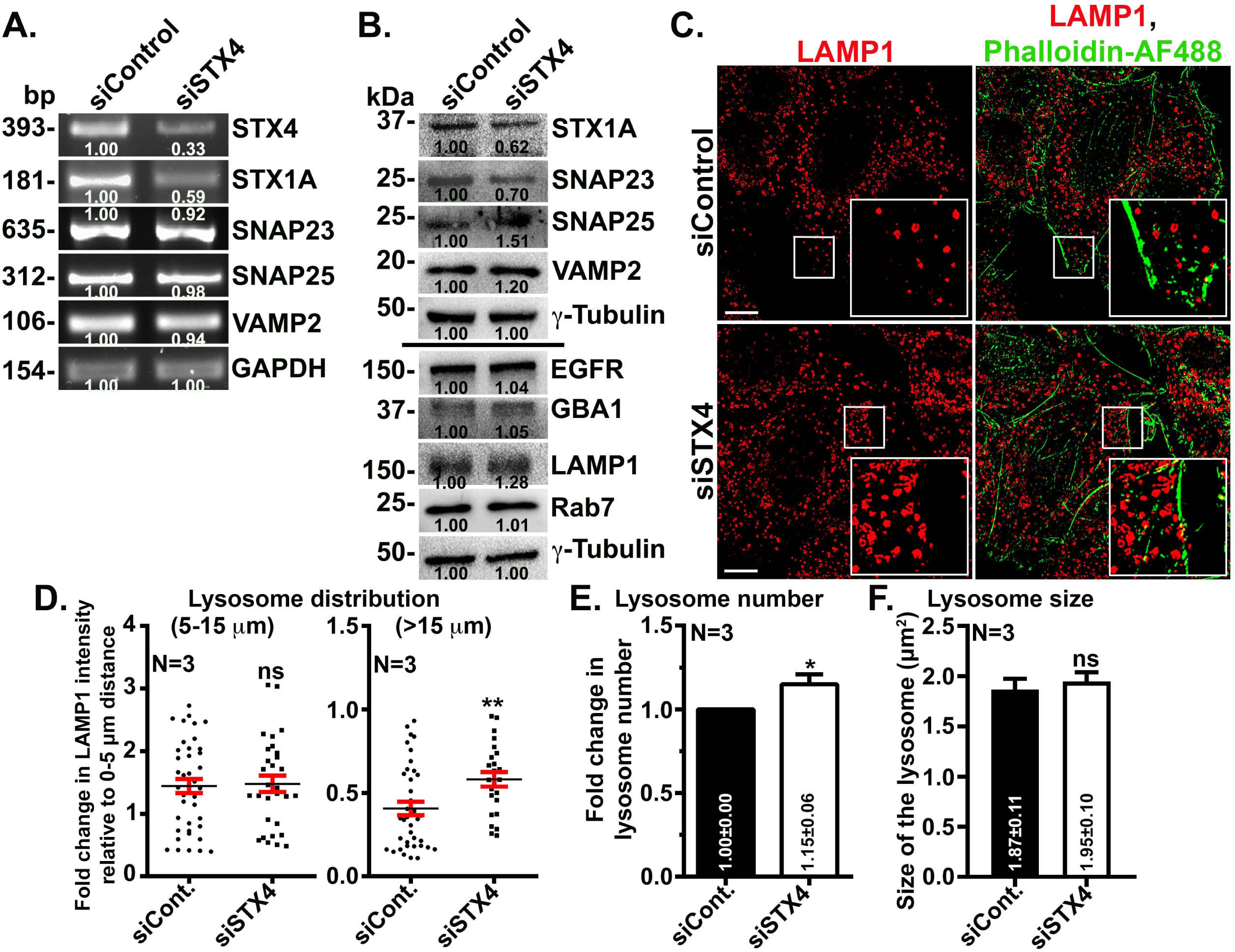
(A) qRT-PCR analysis of different SNAREs in the cell lysates of control and STX4 knockdown cells. The fold change in band intensities was normalized with GAPDH and indicated on the gels. (B) Immunoblotting analysis of control and STX4 knockdown cell lysates. The blots were probed for checking the expression of different SNAREs and lysosome associated and degradative proteins. γ-tubulin is used as the loading control in all immunoblots. The fold change in band intensities was normalized with internal control and indicated on the blots. (C) IFM analysis of HeLa cells transfected with control or STX4 siRNAs. Cells were fixed and stained with anti-LAMP1 antibody and phalloidin-Alexa Fluor 488. Individual and merged panels are shown separately. Insets are magnified views of the white boxed areas. Scale bars, 10 µm. (D) Plots represent the ratio of change in radial profile intensity of lysosomes between 5-15 µm or more than 15 µm distance with respect to 0-5 µm distance, respectively, as a measure of lysosome dispersion in the control and STX4 depleted cells. (E) Plot represents the fold change in lysosome number between control and STX4 depleted cells. (D) Plot represents the size of lysosomes (μm^2^) in control and STX4 depleted cells. The average values in mean±s.e.m. are indicated on the graphs. N=3. ns-non-significant and **p≤0.01.

### Supplementary Videos

**Video S1.** The live-cell TIRF imaging of siControl HeLa cells expressing LAMP1-GFP. Cells were imaged for approximately 2 min.

**Video S2.** The live-cell TIRF imaging of siSTX1A HeLa cells expressing LAMP1-GFP. Cells were imaged for approximately 2 min.

## Notes

### Competing Interest Statement

The authors have declared no competing interest.

